# Fidelity varies in the symbiosis between a gutless marine worm and its microbial consortium

**DOI:** 10.1101/2021.01.30.428904

**Authors:** Yui Sato, Juliane Wippler, Cecilia Wentrup, Rebecca Ansorge, Miriam Sadowski, Harald Gruber-Vodicka, Nicole Dubilier, Manuel Kleiner

## Abstract

In obligate symbioses, partner fidelity plays a central role in maintaining the association over evolutionary time. Fidelity has been well studied in hosts with only a few symbionts, but little is known about how fidelity is maintained in obligate associations with multiple co-occurring symbionts. Here, we show that partner fidelity varies from strict to absent in a gutless marine annelid and its consortium of co-occurring symbionts that provide it with nutrition. We sequenced the metagenomes of 80 *Olavius algarvensis* individuals from the Mediterranean, and compared host mitochondrial and symbiont phylogenies based on single nucleotide polymorphisms across genomes, using a low-coverage sequencing approach that has not yet been applied to microbial community analyses. Fidelity was strongest for the two chemoautotrophic, sulphur-oxidizing symbionts that dominated the microbial consortium in all host individuals. In contrast, fidelity was only intermediate to absent in the sulphate-reducing and spirochaetal symbionts, which occurred in lower abundance and were not always present in all host individuals. We propose that variable degrees of fidelity are advantageous for these hosts by allowing the faithful transmission of their nutritionally most important symbionts and flexibility in the acquisition of other symbionts that promote ecological plasticity in the acquisition of environmental resources.

## Introduction

Beneficial associations between eukaryotic hosts and bacterial partners are ubiquitous, but how these persist stably over evolutionary time remains a source of debate^1-3^. One of the factors that plays a central role in maintaining beneficial symbioses is partner fidelity, defined as the stability of the association between a host and its symbiont over multiple host generations^4^ (see Box 1). In associations with strict fidelity, genetic variants of hosts and symbionts show phylogenetic concordance. Strict fidelity is favoured in associations in which the symbionts are transmitted vertically, that is directly from hosts to their offspring. However, fidelity in vertically transmitted symbioses can be disrupted by host switching, symbiont displacement, acquisition of novel symbionts from free-living microorganisms and symbiont loss. Fidelity is generally weaker in associations with horizontal symbiont transmission in which the symbionts are acquired from free-living microbial populations or co-occurring hosts^4,5^. However, strong fidelity can also occur in symbioses with horizontal transmission if genotype-dependent partner choice ensures the faithfulness of the association^4^.

### Box 1: Definition of key terms used in this study.

As many of the terms below are used inconsistently in the literature, we explain here how we interpret them.

**Vertical transmission:** The direct transmission of a symbiont from a parent to its offspring. In most symbioses the transmission is from mother to offspring (maternal), but there are cases of paternal transmission^6,7^.

**Horizontal transmission:** The transmission of a symbiont to a host from the environment or a co-occurring host^6^.

**Mixed**-**mode transmission:** The transmission of a symbiont by vertical transmission mixed with occasional or frequent events of horizontal transmission over evolutionary time^5^. Note that the transmission of a symbiont community from one generation to the next in which some members are transmitted vertically and others horizontally is not meant here when using this term.

**Partner fidelity:** The stability of the association between host and symbiont genotypes over multiple host generations^8^. Partner fidelity is generated by vertical symbiont transmission, or genotype-dependent partner choice in horizontal symbiont transmission^4^. In this study, we used congruent phylogenies of host mitochondrial genomes and symbiont genomes as an indicator of partner fidelity.

**Partner choice:** The ability of hosts, symbionts or both to preferentially choose their partner. Partner choice describes interactions between individual partners within their lifetime, and is distinct from partner fidelity requiring repeated interactions over evolutionary time^8,9^.

**Partner specificity:** The taxonomic range of partners in an association^10^. Symbiont specificity is defined as the range of symbionts with which a host associates, while host specificity is defined as the range of hosts with which a symbiont associates. In this study, we distinguish partner specificity from partner fidelity, as the former measures the possible diversity of host-symbiont associations, but not the stability of each association.

**Co-inheritance:** The transmission of two or more traits from a host parent to its offspring. Traits can include any combination of phenotypes, genes, alleles, organelles and symbionts. In this study, we use the term to describe the co-inheritance of mitochondria and symbionts from parents to their offspring.

**Phylosymbiosis:** Microbial community relationships that recapitulate the phylogeny of their hosts^11^. Phylosymbiosis tests how similar the composition of microbial communities is to the phylogeny of host species, and can arise through ecological or evolutionary forces. Phylosymbiosis differs from partner fidelity in that the structure of the microbial community is analysed, not the phylogeny of each symbiont taxon.

**Microevolution:** Evolutionary change in a population over short time scales, and generally applied to evolution within a species or conspecific populations.

**Macroevolution:** Evolutionary change over longer time scales, and generally applied to evolution across species and higher taxonomic groups.

Partner fidelity has been well studied in obligate associations with only one or a few symbionts, such as aphids^12-15^, tsetse flies^16^, *Riftia* tube worms^17^, Vesicomyidae clams^18-20^, *Solemya* clams^21^, and the Hawaiian bobtail squid^22^. However, as the number and diversity of symbiont species in a host increases, revealing the patterns of partner fidelity over multiple host generations becomes more challenging^4^. In hosts with highly diverse microbiota, such as corals^23,24^, insects^25-27^ and mammals^28-30^, phylogenetic analyses of microbial marker genes or parts of them, like the 16S rRNA gene, have been used to examine if the microbiome composition reflects the phylogeny of related host species, a pattern termed phylosymbiosis^25^. For a few insect and mammal host taxa, metagenomic analyses have allowed a higher resolution of host and microbiome genetic diversity and revealed phylosymbiosis at shorter evolutionary time scales within species^26,28,31-33^. However, phylosymbiosis is different from partner fidelity in that the structure of the microbial community is analysed, not the co-inheritance of symbiont and host genotypes (see Box 1). Identifying partner fidelity in hosts with highly diverse microbial communities is a challenge because the microbiota generally consist of hundreds to thousands of species and strains that can evolve rapidly and are in continual flux^34^. Well-characterized associations with a limited number and diversity of co-occurring symbionts are ideal for understanding how selective pressure affects different members of a symbiotic community. However, partner fidelity in a symbiosis with multiple, co-occurring symbionts has, to our knowledge, not yet been studied.

Gutless marine annelids offer a unique opportunity for studying partner fidelity in multi-member symbiont communities (Figure 1a). These worms from the subfamily Phallodrilinae (Clitellata, Naididae, genera *Olavius* and *Inanidrilus; sensu* Erséus *et al*.^35^) are regularly associated with at least three to seven symbiont species from different genera and phyla that co-occur within single host individuals^36-43^. The symbionts are harboured in an extracellular region immediately under the outer cuticle of the host (Figure 1b). Over the estimated 50 million years these hosts have evolved from their gut-bearing ancestors^44^, they have become so fully dependent on their symbionts for both nutrition and recycling of their waste compounds that they no longer have a mouth, digestive tract and excretory system^36-38,45^. Symbiont transmission occurs vertically through smearing when the parents deposit their eggs in the sediment, based on transmission electron microscopy and fluorescence *in situ* hybridization (FISH) studies of three host species (*Inanidrilus leukodermatus, Olavius planus*, and *Olavius algarvensis*)^43,46-48^. However, given the morphological similarity of several members of the bacterial symbiont community, these studies could not resolve if all symbionts are inherited through egg-smearing, or if some are acquired horizontally from the environment. Furthermore, there is evidence for host-switching and displacement in the dominant, sulphur-oxidizing symbiont *Candidatus* Thiosymbion over longer evolutionary time^49,50^.

**Figure 1.**
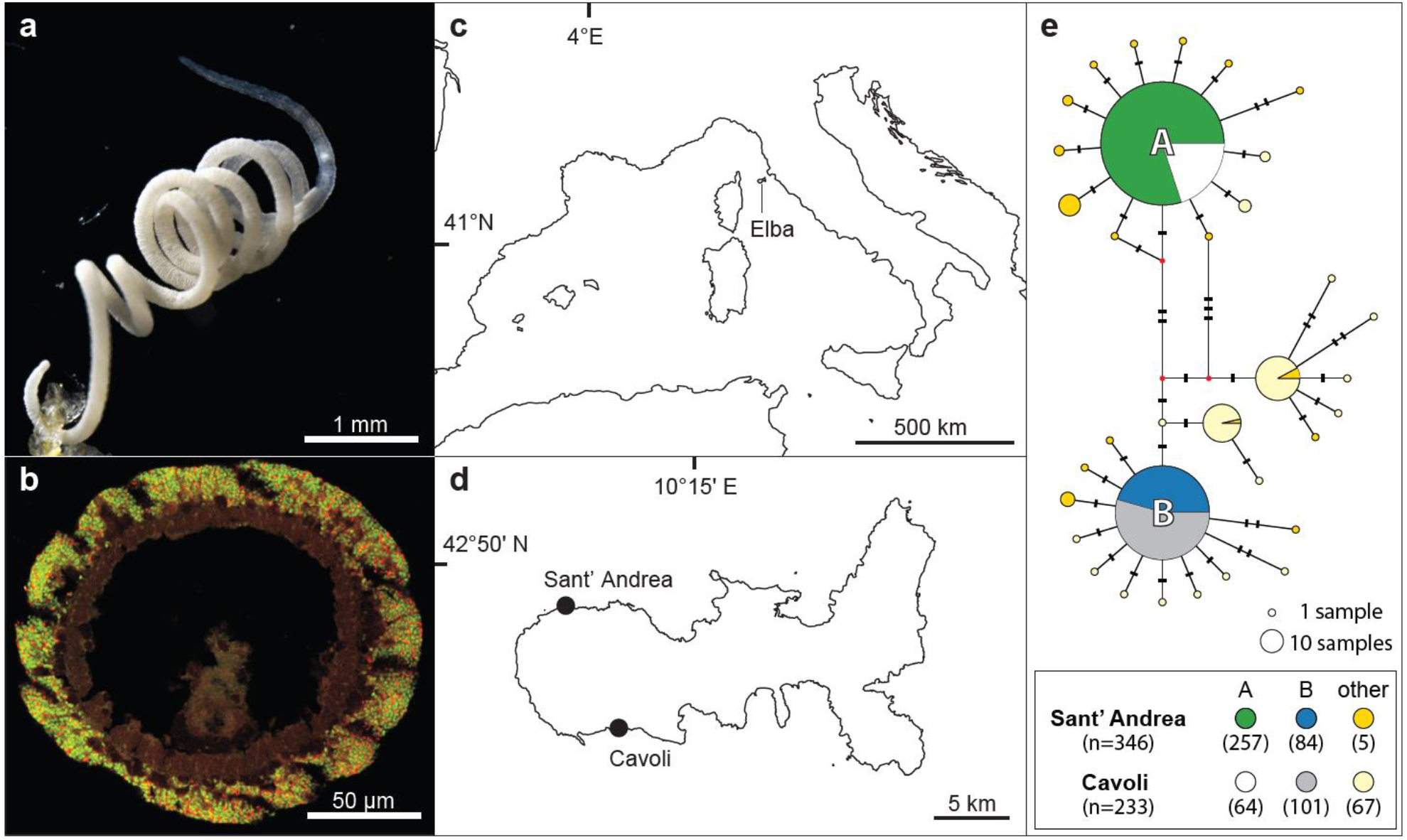
The *O. algarvensis* population in two bays off the island of Elba was dominated by two mitochondrial haplotypes. (**a**) Light microscopy image of *Olavius algarvensis*, (**b**) Fluorescence *in situ* hybridization image of an *O. algarvensis* cross section, highlighting the symbionts just below the cuticle of the host (gammaproteobacterial symbionts in green and deltaproteobacterial symbionts in red, using general probes for these two phyla). Reproduced with permission from Kleiner *et al*.^53^. (**c, d**) Location of the two collection sites, Sant’ Andrea and Cavoli, two bays off the island of Elba in the Mediterranean. (**e**) Haplotype network of mitochondrial cytochrome c oxidase subunit I gene sequences of O. algarvensis individuals from the two collection sites. The two dominant haplotypes A and B co-occurred in both bays. The size of the pie charts corresponds to haplotype frequencies. Hatch marks correspond to the number of point mutations between haplotypes. Nodes depicted by small red points indicate unobserved intermediates predicted by the algorithm in the haplotype network software. The number of individuals identified as haplotype A or B in each bay are in parentheses in the box below the network.

In the best-studied gutless marine annelid, *O. algarvensis*, seven symbiont species have been identified – two sulphur-oxidizing Gammaproteobacteria: *Ca*. Thiosymbion and Gamma3; four sulphate-reducing Deltaproteobacteria: Delta1a, Delta1b, Delta3 and Delta4; and a spirochete^36-39,51,52^. Of these seven symbionts, not all consistently co-occur in all host individuals, but all hosts harbour the sulphur-oxidizing symbiont *Ca*. Thiosymbion, the sulphate-reducing Delta1a or Delta1b symbiont, and the spirochete^39^. The gammaproteobacterial, and possibly the deltaproteobacterial symbionts, autotrophically fix CO_2_, and engage in syntrophic cycling of oxidized and reduced sulphur compounds^36,37,53^. Nutrient transfer to the host occurs via phagolysosomal digestion of the symbionts in the epidermal cells underneath the symbiont layer^54,55^. Evidence from genomic, proteomic and stable isotope analyses indicate resource partitioning, with different symbiont species favouring different energy and carbon sources^37,38,45,54^. This metabolic niche differentiation among these co-occurring symbionts, together with the variability in their abundances across host individuals, indicate different levels of selective pressure, which could be reflected in varying degrees of partner fidelity^56^.

In this study, we examined partner fidelity in populations of *O. algarvensis* from two bays of the island of Elba, Italy, by analysing single nucleotide polymorphisms (SNPs) across 80 metagenomes. For the hosts, we analyzed the phylogeny of their mitochondrial genomes as indicators of vertical symbiont transmission. We then compared mitochondrial phylogeny with that of each symbiont to identify levels of congruence, and correspondingly fidelity, within the microbial consortium. We used a low-coverage sequencing approach for non-model organisms that has not yet been applied to host-microbe associations^57,58^, allowing the analysis of the entire microbial consortium, including low-abundance symbionts.

## Results

### Two mitochondrial haplotypes dominated the host population

To identify mitochondrial genetic diversity in the *O. algarvensis* population, we sequenced the mitochondrial cytochrome c oxidase subunit I (COI) gene of 579 *O. algarvensis* individuals collected over four years from two bays approximately 16 km apart on the island of Elba, Italy (Sant’ Andrea and Cavoli; Figure 1c and 1d). A haplotype network, based on 579 COI sequences of 525 bp, revealed two dominant mitochondrial COI-haplotypes, here termed A and B, that co-occurred at both locations and were distinct from each other by five nucleotides (Figure 1e). We sequenced the metagenomes of 20 individuals from the two dominant A and B haplotypes from both locations (in total 80 metagenomes), and assembled complete circular genomes of host mitochondria (mtDNA) from an arbitrarily-selected metagenome for A- and B-haplotypes. The mtDNAs of A- and B-haplotypes shared 99.3% average nucleotide identity (ANI), and encoded 13 protein-coding genes, 2 ribosomal RNAs, and 21 transfer RNAs (A = 15,715 bp and B = 15,730 bp). The close phylogenetic relatedness between these two host mitochondrial haplotypes that co-occurred in both bays allowed us to examine the effects of both genetics and geographic location on partner fidelity in *O. algarvensis*.

### Symbiont strain diversity was low within host individuals

We assembled metagenome-assembled genomes (MAGs) for each of the seven symbionts in *O. algarvensis* and used them as references for SNP-based analyses (Supplementary Table S1; Supplementary Figures S1 and S2). Symbiont strain diversity within single host individuals was low based on SNP densities in the seven symbiont genomes (0.02 – 0.49 SNP/kbp; Supplementary Table S2b). These values are lower than or comparable to SNP densities reported for vertically-transmitted endosymbionts in the shallow water bivalve *Solemya velum* (0.1 – 1 SNP/kbp)^21^, and considerably lower than SNP densities in horizontally-transmitted symbionts of the giant tubeworm *Riftia pachyptila* (2.9 SNP/kbp)^59^ and deep-sea *Bathymodiolus* mussels (5 – 11 SNP/kbp)^60^. Since strain diversity was low within host individuals, we treated each symbiont within an *O. algarvensis* individual as a single genotype in the analyses described below.

### Sulphur-oxidizing, sulphate-reducing, and spirochete symbionts were found in all host individuals

We assessed the relative abundance of symbionts within each of the 80 *O. algarvensis* individuals by quantifying sequencing read abundances for single-copy genes specific to each symbiont species (Figure 2; Supplementary text 1.1; 162 to 431 single-copy genes per symbiont species). The sulphur-oxidizing symbionts, *Ca*. Thiosymbion and Gamma3, as well as the spirochete, were present in all individuals (Figure 2a). All host individuals also always had sulphate-reducing symbionts (Delta1a, Delta1b, Delta3 and Delta4), but these varied across host individuals and no individuals hosted all of them. Delta3 was detected in only six host individuals, making statistical tests meaningless, and therefore excluded from subsequent phylogenetic analyses.

**Figure 2.**
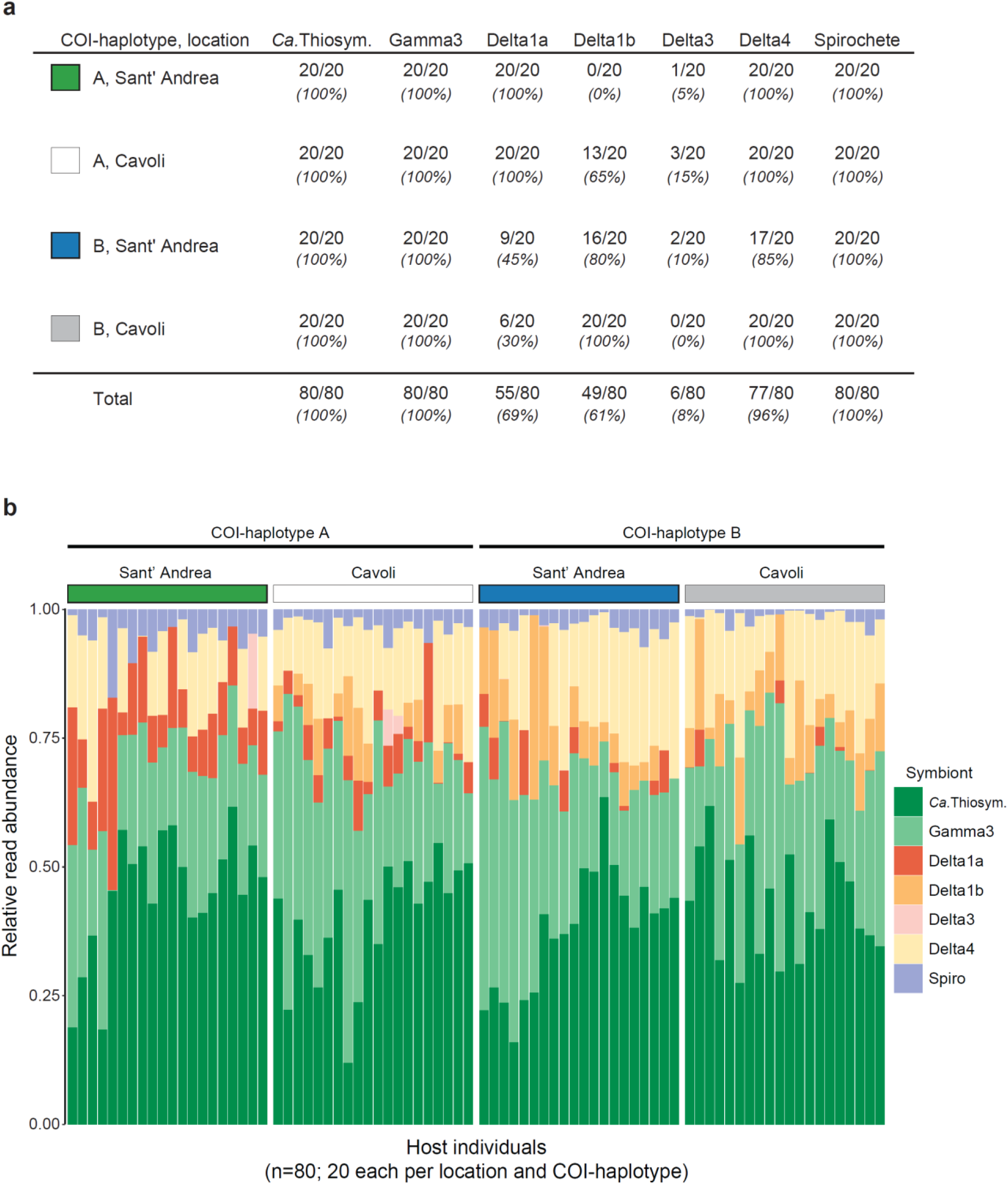
The composition of the symbiont community in 80 *O. algarvensis* individuals. (**a**) The number of *O. algarvensis* host individuals from two mitochondrial lineages (COI-haplotypes A and B) and two locations (Sant’ Andrea and Cavoli) in which the respective symbiont species was detected (n = 80 in total; 20 replicates per location and COI-haplotype; Supplementary text 1.1). (**b**) Relative read abundances of symbionts in the 80 *O. algarvensis* individuals. Each column shows the reads from a single host individual. The sulphur-oxidizing symbionts *Ca*. Thiosymbion (*Ca*. Thiosym.) and Gamma 3 were the most abundant across host individuals, while the abundances of the sulphate-reducing symbionts (Delta1a, Delta1b, Delta3, Delta4) and the spirochete symbiont (Spiro) were consistently lower. Relative abundances of each symbiont were estimated based on metagenomic sequencing reads that mapped to the single-copy genes of each symbiont (Supplementary text 1.1). Relative symbiont abundances based on 16S rRNA gene sequences in the metagenomes were similar (Supplementary text 1.2; Supplementary Figure S4).

Based on relative read abundances mapped to single-copy genes, *Ca*. Thiosymbion was the most abundant symbiont across host individuals (41.6 ± 11.6%; mean ± standard deviation; Figure 2b), while the abundance of Gamma3 symbiont reads varied considerably in host individuals from 0.1 to 61.2% (28.3 ± 11.3%). The summed relative abundances of the sulphur-oxidizing gammaproteobacteria (69.9 ± 7.4%) and sulphate-reducing deltaproteobacteria (26.7 ± 7.2%) showed consistent ratios across the 80 host individuals, regardless of the location, host COI-haplotype, or the combination of these two factors (Supplementary Table S3). Read abundances of the spirochete symbiont were consistently low in host individuals (3.4 ± 2.7%), with one exception of 17.3%.

### Probabilistic SNP-identification increased the number of hosts and SNP-sites for phylogenetic reconstruction of low-abundance symbionts

Conventionally, SNPs in symbiont genomes are identified using a deterministic genotype-calling approach that requires a minimum read-coverage (e.g. at least five-fold^61^; Supplementary text 1.4). In our dataset, this coverage requirement excluded symbionts that occurred at low relative abundances (Supplementary Table 4a). In addition, the deterministic genotype-calling limits the number of available SNP-sites for phylogenetic inference, in our case to a maximum symbiont SNP density of 0.102 SNP/kbp (ranging from 0.006 to 0.102 SNP/kbp; Supplementary Table 4b). To circumvent these limitations, we calculated genotype probabilities and inferred genetic distances from the probability matrices for phylogenetic reconstruction. This allowed us to reconstruct phylogenies of symbionts from more *O. algarvensis* individuals, up to twice as many individuals than using the deterministic approach (between 3% and 131% increase; Supplementary Table S4a). Moreover, the probabilistic approach increased the robustness of our phylogenetic analyses, as it led to a substantial increase in SNP-sites (ranging from 0.035 to 0.752 SNP/kbp; Supplementary Table S4b; see Supplementary text 1.4). Our comparison of phylogenies based on deterministic and probabilistic SNP-identification showed consistent phylogenetic clustering for the samples that could be analysed using both methods (Figure 3; Supplementary Figure S5; Supplementary text 1.4). Therefore, we based our analyses on the probabilistic approach that allowed us to (i) include significantly more samples and (ii) identify more SNP-sites to infer more robust phylogenies (Supplementary Table S4).

**Figure 3.**
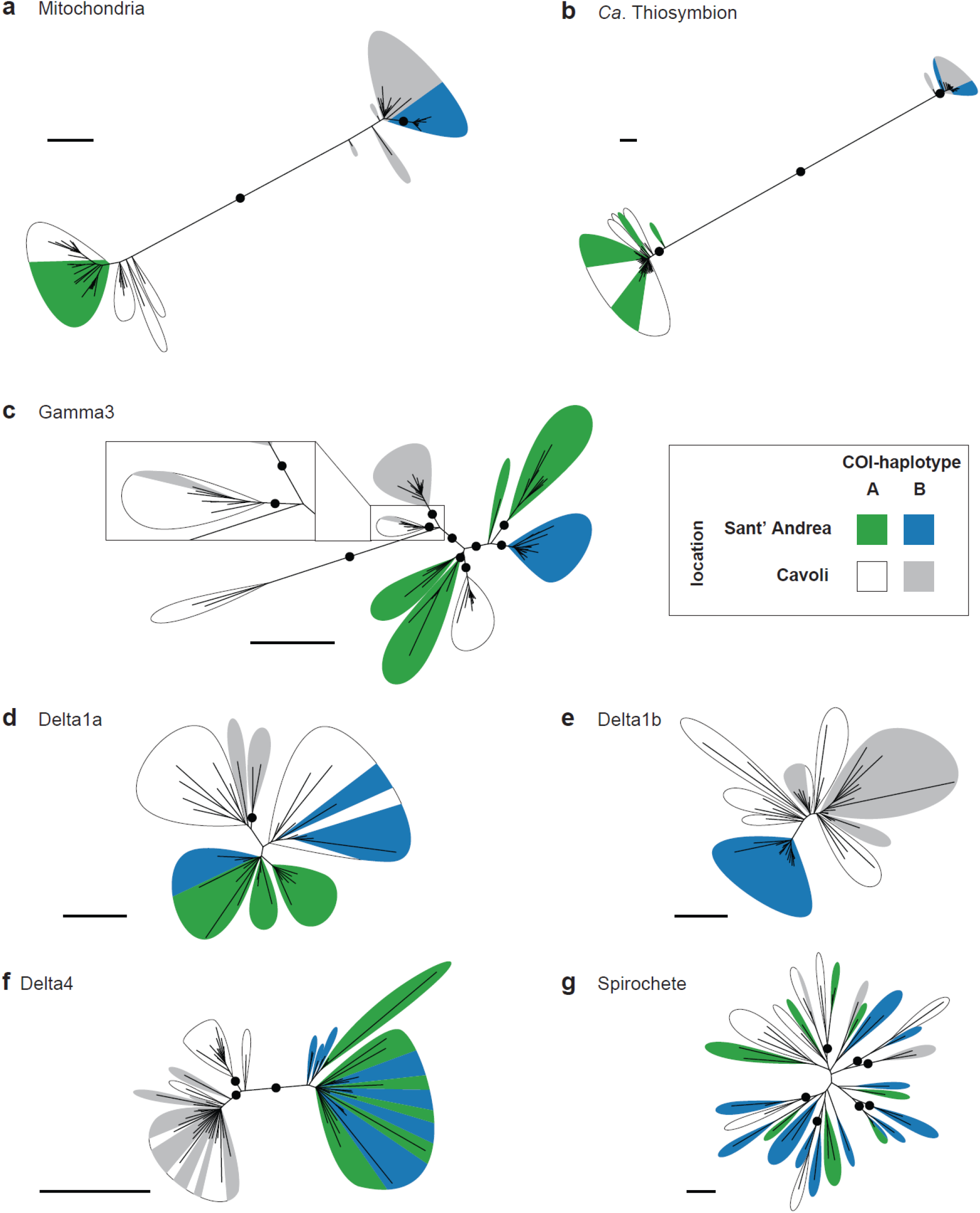
Comparative phylogenetic analyses of *O. algarvensis* and its microbial consortium members revealed variable patterns of congruence across the six symbionts. Phylogenetic trees based on SNPs across genomes of (**a**) host mitochondria (166 SNPs, n = 80), (**b**) *Ca*. Thiosymbion (2872 SNPs, n = 80), (**c**) Gamma3 symbiont (618 SNPs, n = 80), (**d**) Delta1a symbiont (375 SNPs, n =37), (**e**) Delta1b symbiont (624 SNPs, n = 46), (**f**) Delta4 symbiont (675 SNPs, n=67), and (**g**) spirochete symbiont (88 SNPs, n=41). Phylogenies were inferred from genetic distances calculated from posterior genotype probabilities. Scale bars indicate 0.1 substitution per SNP-site. Bootstrap support values >95% are shown in black circles. Supports for branches internal to each coloured leaf were omitted for visibility.

### Congruence of symbiont and mitochondrial phylogenies varied from high to absent

To examine partner fidelity in the *O. algarvensis* symbiosis, we compared the phylogenies of the six most widespread symbionts with that of their hosts’ mitochondrial genomes. The degree of congruence between symbiont and host phylogenies reflects the degree of fidelity between partners, with high congruence indicating strong fidelity and vice-versa^12,13,21^. For the host population, their mtDNA phylogeny revealed a clear divergence between two mitochondrial lineages (termed A- and B-hosts), corresponding to the two COI-haplotypes A and B (Figure 3a). In contrast, the two locations appeared to play a smaller role in shaping mitochondrial phylogeny, with only B-hosts from Sant’ Andrea forming a well-supported clade. However, the phylogenetic relationships within the mitochondrial lineages could not be fully resolved due to their limited genetic divergence (Supplementary Figure S6).

For the six symbionts of *O. algarvensis*, we found marked differences in the congruence of their phylogenies with that of their hosts’ mitochondrial genomes (Figure 3b-g). Congruence was highest in *Ca*. Thiosymbion, which mirrored the mitochondrial phylogeny of their hosts, with symbionts from A- and B-hosts falling into two separate clades (Figure 3b). Phylogenetic relationships within each *Ca*. Thiosymbion clade could not, however, be resolved due to their limited genetic divergence (Supplementary Figure S6b). The Gamma3 symbionts formed nine groups, most of which were statistically supported clades, with each group containing symbionts from either A- or B hosts (Figure 3c). The exception was a Gamma3 symbiont of a B-host from Cavoli that fell in a clade of A-host symbionts from Cavoli (magnified panel in Figure 3c). Location also appeared to affect the phylogeny of Gamma3 symbionts, with symbionts from the same bay forming distinct groups.

For all other symbionts besides the *Ca*. Thiosymbion and Gamma3, there was no phylogenetic divergence between symbionts from A- and B-hosts (Figure 3d-g). Location affected the phylogeny of the Delta4 symbionts, with two well supported clades separating symbionts from Cavoli and Sant’ Andrea (Figure 3f). The Delta1b symbionts also clustered based on their location, but the clades were not statistically supported (Figure 3e). For the Delta1a and spirochete, neither host mitochondrial lineage nor location affected their phylogenetic clustering (Figure 3d-e, 3g).

We further examined congruence between the phylogenies of *O. algarvensis* mitochondria and its symbionts using two additional approaches. First, we compared the pairwise genetic distances of hosts and symbionts using three categories: within A- or B-hosts from the same location (within), between A-and B-hosts (mito), and between the two locations Sant’ Andrea and Cavoli (location). We tested if genetic distances were explained by host mitochondrial lineage or by location, by analysing pairwise genetic distances in ‘within’ vs. ‘mito’, and ‘within’ vs. ‘location’ (Figure 4, Table 1, Supplementary Table S5). For the host, both the mitochondrial lineage and the location had a significant effect on genetic distances, as observed in our phylogenetic SNP analyses. For the symbionts, there was a significant effect of the mitochondrial lineage on genetic distances in *Ca*. Thiosymbion and Gamma3 symbionts, while the effect of location was well supported for the Gamma3 and Delta4 symbionts, again confirming our phylogenetic analyses. An effect of location on genetic distances was also significant for the Delta1b symbionts, but only for B-host symbionts, as they were not detected in A-hosts from Sant’ Andrea.

**Figure 4.**
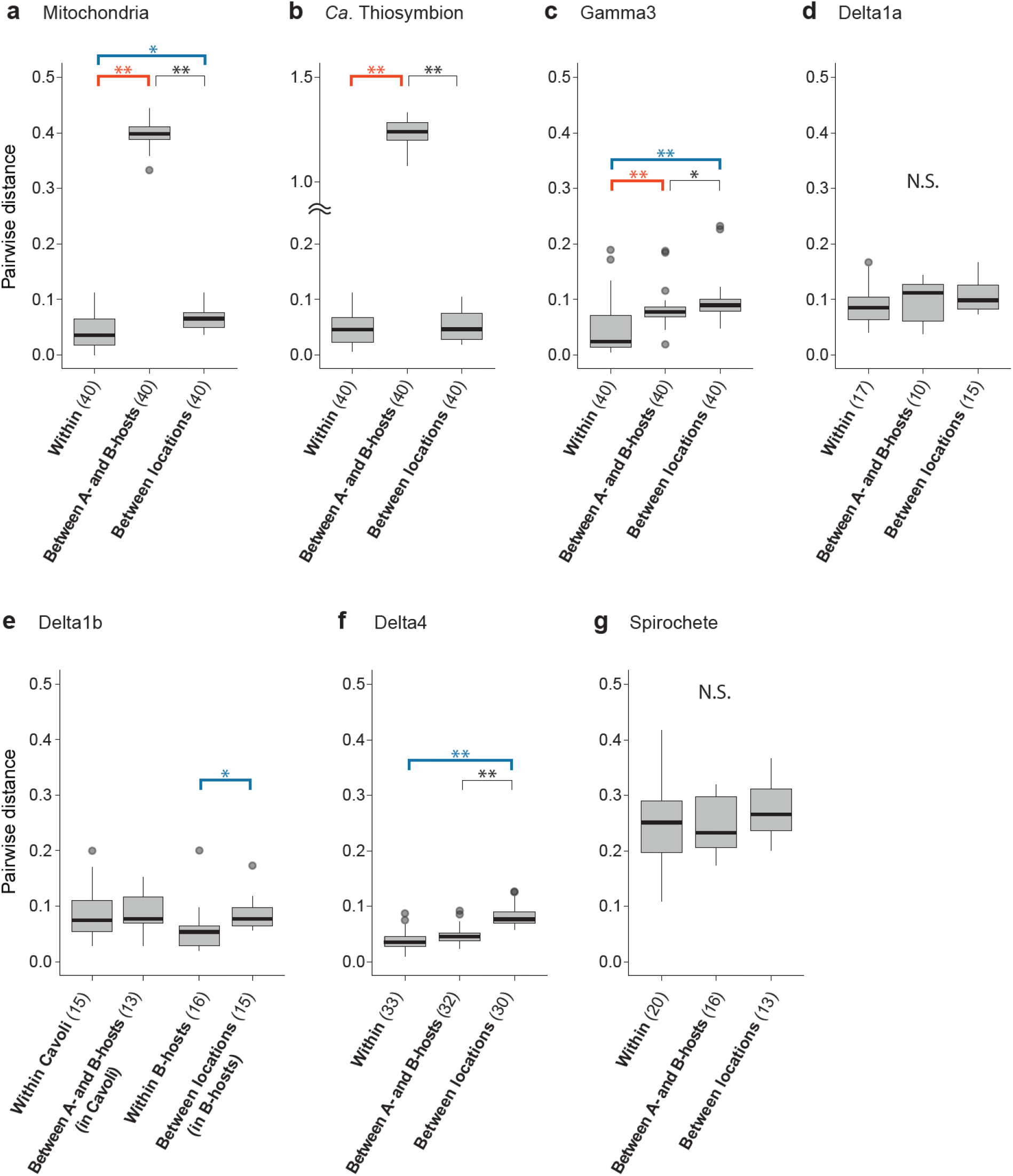
Host mitochondrial lineage and geographic location had a significant effect on the genetic divergence of some but not all symbionts. Host mitochondrial lineages explained genetic divergence in *Ca*. Thiosymbion and Gamma3, while geographic location explained divergence in the Gamma3, Delta1b and Delta4 symbionts. Pairwise genetic distances in *O. algarvensis* mitochondrial genomes and symbionts were calculated from pairs of *O. algarvensis* individuals within the same combination of host lineage (A- or B-host) and location (**Within**), between individuals of A- and B hosts from the same location (**Between A- and B-hosts**), and between individuals from the two locations Sant’ Andrea and Cavoli, but from the same host lineage (**Between locations**). Pairwise genetic distances were compared among these three categories for (**a**) mitochondria, (**b**) *Ca*. Thiosymbion, (**c**) Gamma3 symbiont, (**d**) Delta1a symbiont, (**e**) Delta1b symbiont, (**f**) Delta4 symbiont, and (**g**) spirochete symbiont. Genetic distances were normalized per SNP site and log-scaled. Thick horizontal lines and grey boxes respectively indicate the median and interquartile range (IQR) of observations. Vertical lines show the IQR ± 1.5 IQR range, and outliers out of this range are shown as circles. Numbers in brackets indicate numbers of pairwise comparisons per category tested. Asterisks respectively denote statistical significance (*; p<0.05, **; p<0.01. see Supplementary Table S5). Orange and blue brackets highlight a significant effect on genetic divergence by the mitochondrial lineage and location, respectively. N.S. indicates no significant differences among categories (p>0.05).

**Table 1:** Levels of partner fidelity between *O. algarvensis* and its symbiotic consortia based on analyses in this study.

|  | Observations |  |  |  |  | Indication |
| --- | --- | --- | --- | --- | --- | --- |
| Symbiont | Significant effect on genetic distance |  | Mitochondria-symbiont co-divergence pattern<sup>3</sup> | Relative abundance<sup>4</sup> | Other | Fidelity |
| Ca. Thiosymbion | Mitochondria<sup>1</sup> |  | Strong | 41.6% | Detected in all host individuals. High genomic divergence between host maternal lineages A and B at both locations | Strict |
| Gamma3 | Mitochondria<sup>1</sup> | Location<sup>2</sup> | Present | 28.3% | Detected in all host individuals. A single switch from an A- to a B-host observed | Strong |
| Delta1a |  |  | Weak | 5.3% | Detected in all A-hosts from both locations, but in <50% of B-hosts from both locations | Moderate |
| Delta1b |  | Location<sup>2</sup> | Present | 6.1% | Not detected in A-hosts from Sant' Andrea, but detected in 65 – 100% of other host individuals | Moderate |
| Delta3 | N.A. | N.A. | N.A. | 0.3% | Detected in only 0 – 15% of host individuals | N.A. |
| Delta4 |  | Location<sup>2</sup> | Weak | 14.9% | Detected in 85 – 100% of host individuals | Weak |
| Spirochete |  |  | Absent | 3.4% | Detected in all host individuals | Absent |

<sup>1</sup>; Pairwise genetic distances of symbionts were explained by their host mitochondrial lineage A or B (A- or B-hosts).
<sup>2</sup>; Pairwise genetic distances of symbionts were explained by locations.
<sup>3</sup>; Refer Supplementary Figure S7 and Supplementary Table S6.
<sup>4</sup>; Mean relative read abundances across all host individuals.
N.A.; Not assessed due to the very low abundance of this symbiont in only six host individuals.

We next examined whether the genetic distance of mtDNA between pairs of *O. algarvensis* individuals corresponded to that of their symbionts. To test for this genetic co-divergence, correlations of the pairwise genetic distances were calculated between mtDNA and each of the six symbionts (Supplementary Figure S7, Supplementary Table S6a). The genetic distance of *Ca*. Thiosymbion symbiont had the strongest positive correlation with mtDNA genetic distances (Table 1, Supplementary Figure S7a, Supplementary Table S6a). For the Gamma3, Delta1a, Delta1b, and Delta4 symbionts, we only observed weak positive correlations (Supplementary Figure S7b-e). Genetic co-divergence between the spirochete symbiont and host mitochondria was not detectable (Supplementary Figure S7f, Supplementary Table S6a). Correlation coefficients (Mantel’s R) calculated above for the six symbionts showed a positive correlation with the symbionts’ relative read abundances (Figure 2b; Supplementary Table S6b). In other words, the higher the relative abundance of a symbiont species in the host individuals, the greater the degree of genetic co-divergence between the symbiont species and *O. algarvensis*.

## Discussion

Our analyses revealed that fidelity between the gutless annelid *O. algarvensis* and its endosymbiotic microbial consortium varied from strict to absent. This variability in partner fidelity likely occurred over a short, microevolutionary period, as we analysed a population of *O. algarvensis* from two very closely related mitochondrial lineages (0.7% divergence) that co-occurred in two bays separated by only 16 km. To our knowledge, all previous studies on the stability of symbioses in hosts that are obligately associated with multiple symbionts have focussed on congruence of host and symbiont phylogenies over macroevolutionary timescales (e.g.^14,62-64^). Our study highlights the importance of examining fidelity over microevolutionary timescales, as it was central to revealing the broad range of fidelity across the members of the *O. algarvensis* symbiont community from strict over intermediate to absent. Over macroevolutionary timescales, strict fidelity is disrupted in *Ca*. Thiosymbion (see below), and it is unlikely that we would have detected the strong to moderate levels of fidelity in other members of the *O. algarvensis* symbiont community over longer evolutionary time.

### Varying degrees of partner fidelity indicate a spectrum of mixed-modes between vertical and horizontal transmission

Different degrees of partner fidelity across the microbial consortium of *O. algarvensis* reflect the faithfulness with which the symbionts are transmitted from one generation to the next. The symbionts of marine gutless annelids are transmitted vertically through egg-smearing, during which the egg passes through symbiont-rich tissues termed genital pads^46-48^. The egg is then encased in a cocoon and deposited in the sediment. This process offers opportunities for horizontal transmission of bacteria from the surrounding sediment or co-occurring hosts. Indeed, previous studies have shown that over macroevolutionary time, host switching and displacement disrupt fidelity in *Ca*. Thiosymbion^49,50^. Our analyses of *O. algarvensis* suggest that on microevolutionary scales, transmission is strictly vertical for *Ca*. Thiosymbion, given the strong phylogenetic congruence and genetic co-divergence between these symbionts and their hosts’ mitochondria, independent of their collection site (Supplementary text 1.6). We also observed strong fidelity in the Gamma3 symbiont, with only a single switching event from an A-to a B-host in Cavoli (Figure 3c). In contrast, partner fidelity is intermediate to absent for the deltaproteobacterial and spirochete symbionts. It is therefore likely that these symbionts are regularly acquired horizontally from a free-living population or other co-occurring host individuals (Supplementary text 1.6). For the Gamma3, Delta1b and Delta4 symbionts, we also observed effect of their geographic location on their genetic distances (Figure 4, Table 1). Explanations for this effect include differences in the geographic distribution of genotypes for these symbionts in the environment as well as isolation-by-distance effects on partner choice, but cannot be resolved without additional in-depth analyses of the free-living symbiont population.

### How can we explain such different degrees of partner fidelity in the *O. algarvensis* symbiosis

Strong partner fidelity is widespread in associations in which the symbionts are critical for the host’s survival and fitness^2,65^. In *O. algarvensis*, the relative abundance of symbionts was positively correlated with fidelity, with the highest fidelity detected for their sulphur-oxidizing symbionts *Ca*. Thiosymbion and Gamma3. These symbionts are the primary producers for their gutless hosts by autotrophically fixing CO_2_ into organic compounds^36,38,45^. All *O. algarvensis* individuals from both bays harboured these symbionts, and they consistently dominated the microbial consortia at 46 – 86%. Given that *O. algarvensis* gains all of its nutrition by digesting its endosymbionts via endocytosis^54,55^, those symbionts that provide most of its nutrition are likely most strongly selected for, as they most directly affect the fitness of their host.

In contrast, selective pressures on maintaining the symbiotic association are likely relaxed for symbionts that are less critical for their host’s fitness or are functionally redundant. The deltaproteobacterial, sulphate-reducing symbionts play an important role by producing reduced sulphur compounds as energy sources for the sulphur-oxidizing bacteria, particularly when these compounds are limiting in the sediment environment^36^. However, all four deltaproteobacterial symbionts reduce sulphate to sulphide^36-38,51,52^, making this trait functionally redundant among these symbionts. Correspondingly, the presence and abundance of the four sulphate-reducing symbionts varied across host individuals, with different combinations of one to three of these symbionts in each host. This pick and mix pattern indicates that *O. algarvensis* fitness is not dependent on any particular deltaproteobacterial symbiont species. The lower relative abundance of reads from deltaproteobacterial symbionts (summed average 27%) compared to those of the sulphur oxidizers (70%), as well as their limited biomass^28,29^, further indicates that the deltaproteobacterial symbionts are not as essential for their host’s nutrition as *Ca*. Thiosymbion and Gamma3.

The absence of fidelity in the spirochete symbionts was unexpected, given their regular presence in *O. algarvensis* individuals and in other gutless annelids from around the world^40,66^ (Supplementary Figure S2). While the metabolism of this symbiont is not yet known, it was by far the least abundant symbiont in *O. algarvensis* in this study (3% relative read abundance) as well as previous ones^37-39,53,54^, indicating a limited role of these symbionts in their hosts’ nutrition.

### Having your cake and eating it too: The advantage of flexibility in partner fidelity

The lack of stringent partner fidelity between *O. algarvensis* and its deltaproteobacterial and spirochete symbionts indicates that these hosts regularly acquire novel intraspecific genotypes from the environment or from co-occurring host individuals. These newly acquired intraspecific genotypes, together with the interspecific variability in the deltaproteobacterial community across host individuals, could expand the ecological niche of *O. algarvensis*, and enable the population to adapt to fluctuating environments. For example, new symbiont genotypes could be better adapted to the environment, provide greater flexibility in the use of resources from the environment, and enable resilience to seasonal and long-term temperature changes^5,67-69^. On the other hand, weak partner fidelity can be costly for the host, including the failure to find suitable symbionts, acquisition of harmful bacteria, or association with ‘cheaters’ that do not provide mutualistic benefits^70^. However, these costs are likely to be minimal for the deltaproteobacterial symbionts, as their partner quality depends largely on their ability to produce reduced sulphur compounds, a trait that is characteristic for all sulphate-reducing bacteria, and for which cheating is not possible.

For *O. algarvensis*, fluctuating environmental conditions may be more critical selective forces than the costs of weak partner fidelity. *O. algarvensis*, like other marine annelids, does not have a pelagic life stage and can likely not disperse as widely as many other infaunal invertebrates^47^. These hosts therefore face the risk of local extinction if environmental conditions become unsuitable for their symbionts. Free-living bacteria rapidly adapt to new environmental conditions^71^, and horizontal acquisition of symbionts from the environment can increase the host’s potential to adapt to environmental challenges rapidly^72^. The benefit of long-term survival via an evolutionary bet-hedging^73,74^ may therefore outweigh the cost of recruiting less-favourable symbionts^56^.

### Probabilistic approaches to SNP-identification expand ecological and evolutionary studies of microbial communities

Metagenomic analyses of intraspecific genetic variability in microbial communities typically rely on deep sequencing of each genotype to obtain sufficient SNPs across their genomes^75^, incurring substantial sequencing costs. In this study, we reconstructed phylogenies of symbionts directly from genotype probabilities across their genomes, rather than calling genotypes. This probabilistic approach accurately identifies SNPs from low-coverage sequencing data by accounting for the uncertainties of genotyping^57^. This approach has several advantages, including the lower costs of low-coverage sequencing, the ability to analyse the genetic diversity of low-abundance species, and an increased robustness of phylogenetic analyses through the recovery of higher SNP numbers. Our study highlights how approaches using genotype probabilities can be applied to the population genetics of host-associated microbiota with variable abundances. Furthermore, our results indicate that probabilistic approaches can also be used to study evolutionary dynamics in free-living microbial communities, thus greatly expanding our toolbox for understanding non-cultivable microorganisms.

### Conclusions

We showed that partner fidelity varies from strict to absent in the association between the gutless marine annelid *O. algarvensis* and its microbial consortium. This variability in fidelity was unexpected given that these hosts transmit their symbionts vertically via egg-smearing. Our results highlight the importance of examining partner fidelity at microevolutionary scales, as over longer evolutionary time, strict vertical transmission is rare in most symbioses^69,76^. Understanding the processes that drive fidelity within associations over short to long evolutionary time will help identify the benefits and costs in maintaining symbiotic associations. Such efforts should ideally encompass increasing geographic and taxonomic scales, beginning with local host populations and expanding to regional intraspecific analyses to large-scale global analyses across host species, e.g.^77-80^. Rapid advances in high-throughput sequencing combined with substantial reductions in sequencing costs using probabilistic SNP-calling now make such studies feasible, and will contribute to revealing the driving forces that shape the complex and fluid nature of multi-member symbioses^68,81-83^.

## Methods

### Specimen collection and host mitochondrial lineage screening

A total of 579 *O. algarvensis* individuals from two locations on Elba, Italy (Sant’ Andrea; 42°48’31”N / 10°08’33”E, and Cavoli, 42°44’05”N / 10°11’12”E; Figure 1c and 1d; n = 346 and 233, respectively) were screened for their mitochondrial lineages based on their mitochondrial COI gene sequences (COI-haplotypes). *O. algarvensis* individuals were collected between 2010 and 2016 from sandy sediments in the vicinity of *Posidonia oceanica* seagrass beds at water depths between 7 and 14 m, as previously described^38^. Live specimens were (i) flash-frozen in liquid nitrogen and stored at -80° C, or (ii) immersed in RNAlater (Thermo Fisher Scientific, Waltham, MA, USA) and stored at 4° C. DNA was individually extracted from single worms using the DNeasy Blood & Tissue Kit (Qiagen, Hilden) according to the manufacturer’s instructions. 670 bp of the COI gene was amplified with PCR using the primer set of COI-1490F (5’-GGT-CAA-CAA-ATC-ATA-AAG-ATA-TTG-G-3’) and COI-2189R (5’-TAA-ACT-TCA-GGG-TGA-CCA-AAA-AAT-CA-3’) as previously described^84^. PCR-amplicons were sequenced using the BigDye Sanger sequencing kit (Life Technologies, Darmstadt, Germany) with the COI-2189R primer on the Applied Biosystems Hitachi capillary sequencer (Applied Biosystems, Waltham, USA) according to the manufacturer’s instructions. COI sequences were quality-filtered with a maximum error rate of 0.5% and aligned using MAFFT v7.45^85^ in Geneious software v11.0.3 (Biomatters, Auckland, New Zealand). A COI-haplotype network was built on a 525 bp core alignment using the TCS statistical parsimony algorithm^86^ implemented in PopART v1.7^87^. Twenty *O. algarvensis* individuals from each ‘host group’ (i.e. the combination of sample location, Sant’ Andrea or Cavoli, and COI-haplotype A or B; 4 groups in total) were randomly selected for metagenomic sequencing (n=80 individuals total).

### Metagenome sequencing

Sequencing libraries were constructed from the DNA extracted from single worms using a Tn5 transposase purification and tagmentation protocol^88^. The Tn5 transposase was provided by the Protein Science Facility at Karolinska Institutet Sci Life Lab (Solna, Sweden). Quantity and quality of DNA samples were checked with the Quantus Fluorometer with the QuantiFluor dsDNA System (Promega Corporation, Madison, WI, USA), the Agilent TapeStation System with the DNA ScreenTape (Agilent Technologies, Santa Clara, CA, USA), and the FEMTO Pulse genomic DNA analysis kit (Advanced Analytical Technologies Inc., Heidelberg, Germany) prior to library construction. Insert template DNA was size-selected for 400 – 500 bp using the AMPure XP (Beckman Coulter, Indianapolis, IN, USA). Paired-end 150 bp sequences were generated using the Illumina HiSeq3000 System (Illumina, San Diego, CA, USA) with an average total yield of 2.5 Gbp per sample (2,548 ± 715 Mbp (mean ± SD)). Construction and quality control of libraries and sequencing were performed at the Max Planck Genome Centre (Cologne, Germany).

### Assembly of reference genomes of endosymbionts and *O. algarvensis* mitochondrion

Metagenome-assembled genomes (MAGs) of the *Ca*. Thiosymbion, Gamma3, Delta4 and spirochete symbionts were *de-novo* assembled from a deeply-sequenced metagenome of an *O. algarvensis* individual available at the European Nucleotide Archive (ENA) project PRJEB28157 (Specimen ID: OalgB6SA; approx. 7 Gbp; Supplementary Table S1). In addition, MAGs of Delta1a, Delta1b and Delta3 were obtained from the public database deposited under the ENA project accession number PRJEB28157^51,52^. For the *de-novo* assembly, raw metagenomic reads were first adapter-trimmed and quality-filtered with length ≥ 36 bp and Phred quality score ≥ 2 using *bbduk* of BBTools v36.86 (http://jgi.doe.gov/data-and-tools/bbtools), and corrected for sequencing errors using BayesHammer^89^ implemented in SPAdes v3.9.1^90^. Clean reads were assembled using MEGAHIT v1.0.6^91^, and symbiont genome bins were identified with MetaBAT v0.26.3^92^. Bins were assigned to the *Ca*. Thiosymbion, Gamma3, Delta4 and spirochete symbionts based on 99 – 100% sequence matches with their reference 16S rRNA gene sequences (NCBI accession numbers AF328856, AJ620496, AJ620497, AJ620502^36,39^, respectively). MAGs were further refined using Bandage v0.08.1^93^ by identifying and inspecting connected contigs on assembly graphs. Completeness and contamination of the MAGs were estimated with CheckM version 1.0.7^94^, and assembly statistics of symbiont genomes were calculated with QUAST v5.0.2^95^ (Supplementary Table S1).

Complete mitochondrial genomes (mtDNA) were assembled from two metagenomes of *O. algarvensis* from Sant’ Andrea representing the two COI-haplotypes A and B (Specimen IDs: “OalgSANT_A04” and “OalgSANT_B04”, respectively). Duplicated sequences due to PCR-amplification during the library preparation were first removed from raw sequences using FastUniq v1.1^96^ prior to adapter-clipping and quality-trimming with Phred quality score ≥ 2 using Trimmomatic v0.36^97^. A preliminary mtDNA scaffold was first generated by iterative mapping of the clean reads to a reference COI sequence of *O. algarvensis* (NCBI accession number KP943854; as a bait sequence) with MITObim v1.9^98^. Mitochondrial reads were then identified by mapping to the mtDNA scaffold using *bbmap* in BBTools, and mtDNA was assembled from the identified reads with SPAdes. A circular mtDNA was identified on assembly graphs in Bandage, and annotated using MITOS2 webserver (http://mitos2.bioinf.uni-leipzig.de) to confirm the completeness^99^.

### Characterization of symbiont community composition

The taxonomic composition of the *O. algarvensis* metagenomes was first screened by identifying 16S rRNA gene sequences using phyloFlash v3.3-beta1^100^ using the SILVA SSU database release 132^101^ as reference. The 16S rRNA gene sequences of *O. algarvensis* symbionts were assembled with SPAdes implemented in phyloFlash, and aligned using MAFFT in Geneious to identify SNP sites. Chimeric and incomplete (<1,100 bp) SSU sequences were identified in the alignment and excluded.

Relative abundances of *O. algarvensis* symbionts were estimated by mapping metagenome reads to a collection of symbiont-specific sequences of single-copy genes extracted from the genome bins. Orthologous single-copy gene sequences were first identified within each of the reference symbiont MAGs with CheckM. To ensure unambiguous taxon-differentiation, duplicated genes detected in each symbiont MAG (i.e. those labelled as ‘contamination’ in CheckM) as well as sequences sharing >90% nucleotide identity between multiple symbiont species (checked with CD-HIT v4.5.4^102^; 8 cases identified between the Delta1a and Delta1b symbionts) were removed from the final reference sequences of single-copy genes. Metagenomic reads (quality-filtered with the same processes as for the mtDNA assembly above) matching to the singe-copy gene sequences were quantified using Kallisto v0.44.0^103^ (Supplementary text 1.1). Symbiont composition estimates were plotted in R using ggplot2 package v3.2.1^104^.

To examine whether our phylogenetic analysis of each *O. algarvensis* symbiont reflects a single dominant genotype per host individual, levels of symbiont genotype diversity within a host individual were assessed by calculating SNP-densities (the number of SNP sites per Kbp of reference genome) with previously established procedures^60^. This analysis was performed using a set of publicly available deeply-sequenced metagenomes of *O. algarvensis* (listed in Supplementary Table S2a) to ensure that SNP-densities were estimated using sufficient symbiont read-coverages and that these estimates could be compared to those in other studies performing similar assessments^21,59,60^.

### Identification of single nucleotide polymorphisms and phylogenetic reconstruction

For identification of SNPs in the genomes of symbionts and mitochondria, quality-controlled reads were first unambiguously split into different symbiont species using *bbsplit* of BBTools, using the reference symbiont MAGs described above. Mitochondrial reads were identified with the reference mtDNA sequence derived from the specimen “OalgSANT_A04” (Sant’ Andrea, COI-haplotype A) using *bbmap* with a minimum nucleotide identity of 95%; This step was performed to remove potential contaminations from sequences of nuclear mitochondrial pseudogenes divergent from the mtDNA^105^, while ensuring successful mapping of mtDNA reads in the analysis for both A- and B-hosts given the high nucleotide identity of mtDNAs between these two lineages (>99%; see results).

To reconstruct phylogenies of symbionts and mitochondria, SNPs in genomes of symbionts and host mitochondria were identified using two approaches; based on (**i**) posterior genotype probabilities without genotype calling, and (**ii**) deterministic genotyping only at genetic positions that were deeply sequenced in all samples. For the SNP-identification without genotyping (**i**), the symbiont- and mitochondrial-reads identified above were mapped onto individual reference genomes of symbionts and mitochondria using *bbmap*. Mapping files were filtered based on mapping quality using samtools v1.3.1^106^ and BamUtil v1.0.14^107^, deduplicated with *MarkDuplicates* of Picard Toolkit v2.9.2 (https://github.com/broadinstitute/picard), and realigned around indels with Genome Analysis Tool Kit v3.7^108^. Posterior genotype probabilities were calculated with ANGSD v0.929^109^. ANGSD is widely used for studies of diploid organisms to infer genotypes from low-coverage sequencing data while taking sequencing errors into account^109-111^. To deal with haploid genotypes in ANGSD, all genotypes were assumed to be ‘homozygous’ by setting an inbreeding coefficient F of ‘1’ and a uniform prior were specified for posterior probability calculation (T.S. Korneliussen, 2018, pers. comm.). SNP sites were identified as reference nucleotide positions that were covered ≥1× by all samples and showed statistically significant support (SNP p-value < 0.01). When no SNP site was found due to very low or lacking reads from a symbiont, these samples were excluded based on a cut-off of lateral coverages (i.e. % reference genetic sites covered by reads; Supplementary Table S4). For symbionts and mitochondria, their pairwise genetic distances in host individuals were obtained from the matrix of genotype probabilities using NGSdist v1.0.2^58^. Phylogenetic trees with bootstrap-support were computed from the resulting distant matrix using FastME v2.1.5.1^112^ and RAxML v8.2.11^113^.

For the SNP-identification by deterministic genotyping (**ii**), the same symbiont- and mitochondrial-reads were analysed with the SNIPPY pipeline v3.2 (https://github.com/tseemann/snippy), with the same reference genomes of mtDNA and symbionts as above (Supplementary text 1.4).

### Analyses of phylogenetic patterns of symbionts based on host mitochondrial lineages and locations

To examine which factors drive the patterns of genetic divergence in mitochondria and symbionts, their pairwise genetic distances were statistically compared in 3 categories of host pairs; (**a**) within the same combination of location plus A- or B-hosts, (**b**) between A- and B-hosts (A vs. B in Cavoli, and A vs. B in Sant’ Andrea), and (**c**) between locations (Cavoli vs. Sant’ Andrea in A-hosts, and Cavoli vs. Sant’ Andrea in B-hosts). Sample pairs in each category were randomly selected without replacement to ensure data independence. For the Delta1b symbionts, we separately compared (**a**) vs. (**b**) within Cavoli, and (**a**) vs. (**c**) within B-hosts, because Delta1b did not occur in A-hosts in Sant’ Andrea. The statistical analyses were performed for each of the symbionts and mitochondria using Kruskal-Wallis rank sum test and Dunn post-hoc tests with p-value adjustment by controlling the false-discovery rate, using R core package v3.4.2^114^ and FSA package v0.8.25^115^. To examine genetic co-divergence patterns between a symbiont and host mtDNA, regressions of pairwise genetic distances estimated with NGSdist were examined using Mantel tests implemented in the R-package *vegan* v2.4.4^116^. A correlation between Mantel’s Rs and relative abundances of symbionts based on reads mapped to single-copy genes was tested with Kendall’s rank correlation tau using the function *test*.*cor* in R’s core package.

## Data and script availability

Raw metagenome sequences, and reference genomes of mitochondria and symbionts generated in this study were deposited in the European Nucleotide Archive (ENA) under accession number PRJEB42310. Annotated bioinformatic scripts with all specific parameters, as well as reference single-copy gene sequences, are available in a GitHub depository (https://github.com/yuisato/Oalg_linkage).

## Acknowledgements

This work was supported by the Max Planck Society, a Gordon and Betty Moore Foundation Marine Microbial Initiative Investigator Award to ND (Grant GBMF3811), a U.S. National Science Foundation award to MK (grant IOS 2003107), the USDA National Institute of Food and Agriculture Hatch project 1014212 (MK), and the European Union’s Horizon 2020 research and innovation program under the Marie Sklodowska-Curie grant agreement No 660280 (CW). The authors thank the Hydra Institute for logistical support during sample collection, and Toby Kiers and members of the Department of Symbiosis for valuable discussions.

## Author Contributions

ND, MK, CW and JW conceived the study and MK, ND, YS, JW and HGV designed the work. YS, JW, CW, MS and MK acquired materials and data, YS, JW and RA analysed data. YS, JW, RA, ND and MK interpreted results. YS wrote the manuscript with support from ND and MK, and revisions from all other co-authors.

