## Supplementary for "Fidelity varies in the symbiosis between a gutless marine worm and its microbial consortium"

**Contents:**

1. **Supplementary text**

   **1.1.** Quantification and detection of symbionts based on single-copy marker genes

   **1.2.** Assessment of symbiont community compositions based on 16S rRNA genes

   **1.3.** Symbiont 16S rRNA gene sequences indicated a linkage between haplotypes of *Candidatus* Thiosymbion and host mitochondria

   **1.4.** Reconstruction of mitochondria and symbiont phylogenies using a deterministic genotyping approach to SNP-identification

   **1.5.** Estimation of the effective population size of symbionts within an *Olavius algarvensis* individual based on genome-wide SNP abundance

   **1.6.** Transmission modes of symbionts in *O. algarvensis*

   **Reference cited in supplementary text**

2. **Supplementary figures**

   **Supplementary Figure S1** Phylogenomic tree of symbionts in *O. algarvensis* in relation to reference bacterial genomes

   **Supplementary Figure S2** Phylogeny of 16S ribosomal RNA gene sequences for symbionts in *O. algarvensis* in relation to reference bacterial sequences

   **Supplementary Figure S3** Relative proportions of mapped reads to each of single-copy genes within a single host per symbiont species

**Supplementary Figure S4** Symbiont composition of individual *O. algarvensis* samples of two COI-haplotypes (A and B) from two locations (Sant' Andrea and Cavoli) based on 16S rRNA gene

**Supplementary Figure S5** Core SNP-trees based on genotypes using a deterministic approach to SNP-identification

**Supplementary Figure S6** Phylogenies of mitochondria and *Candidatus* Thiosymbion within the major mitochondrial lineages A and B

**Supplementary Figure S7** Correlation between mitochondrial pairwise genetic distance and pairwise genetic distance of symbionts

**Supplementary Figure S8** Effective population size estimates of the symbiont per *O. algarvensis* individual

**Supplementary Figure S9** Sequence alignments of 16S ribosomal RNA genes of *O. algarvensis* symbionts

#### 3. Supplementary tables

**Supplementary Table S1** Summary statistics of the reference metagenome-assembled genomes of *O. algarvensis* symbionts

**Supplementary Table S2** Assessment of strain diversity of symbionts within *O. algarvensis* individuals

**Supplementary Table S3** Statistical comparisons of ratios between summed relative abundances of deltaproteobacterial and gammaproteobacterial symbionts

**Supplementary Table S4** Comparison of SNP-identification methods

**Supplementary Table S5** Statistical comparisons of pairwise genetic distances for the host mitochondria and symbionts

**Supplementary Table S6** Statistical tests on genetic co-divergence patterns of host mitochondria and symbionts

### 1. Supplementary text to Results and Discussion

##### 1.1 Quantification and detection of symbionts based on single-copy marker genes

In the 80 metagenomes of *Olavius algarvensis*, we assessed (i) symbiont prevalence (the number of host individuals in which the respective symbiont species was detected; n = 20 hosts per host group) and (ii) relative abundances of the symbionts. For this we quantified sequences of single-copy genes (SCGs) that are specific to symbiont species. Between 162 and 431 SCGs per symbiont species were extracted from their reference genomes (Supplementary Figure S3).

Upon assessing the distribution of read coverages of individual SCGs within each symbiont, we identified a small subset of SCGs that indicated particularly high read coverages, especially in the *Candidatus* (*Ca.*) Thiosymbion symbiont (Supplementary Figure S3). Metagenomes with insufficient reads aligned to SCGs (mean TMP counts < 5) were excluded from this read-distribution assessment to ensure that all SCGs of the target symbiont were covered with reads. A closer look at sequence alignment of highly mapped SCGs revealed that these SCGs contain short repeat sequences that attracted high abundances of reads mapped erroneously. Further inspection of the sequence alignment indicated that similar short repeat sequences were also present in the corresponding symbiont genome, which likely resulted in the overestimation of the small number of SCGs. To exclude these SCGs from the symbiont abundance calculation without causing erroneous underestimation, we first ordered SCGs by read coverage within each sample. We then calculated the mean read coverage from SCGs ranked between 25 and 75 percentiles of the order (i.e. genes in the interquartile range; including SCGs with no mapped reads in the ranking). Although we obtained a robust estimate for symbiont relative abundance using this approach, the quantification of a particular symbiont required its metagenomic sequences mapping to more than 25% of the reference SCGs (i.e. between 41 and 108 genes depending on the symbiont). This would have set the limit of symbiont detection on the conservative side. To better control false negatives and false positives in symbiont detection, we set the criteria for the detection of a symbiont as (i) read-mapping on at least 15% of its reference SCGs (i.e. between 24 and 65 genes depending on the symbiont), as well as (ii) the detection of the reads matching its 16S ribosomal RNA gene (SSU), given that the SSU sequence is the well-established taxonomic marker used for bacterial identification[1]. Results of symbiont detection are summarised in Figure 2a.

#### 1.2. Assessment of symbiont community compositions based on 16S ribosomal RNA genes

In addition to the SCG-based approach to symbiont quantification above, relative abundances of *O. algarvensis* symbionts were estimated also by mapping metagenome reads to reference SSU sequences of the symbionts. For SSU-based estimation, quality-filtered reads matching with symbiont SSU sequences were quantified with Kallisto, using representative SSU reference sequences (NCBI accession numbers: AJ620502 (spirochete), AJ620497 (Delta4), AJ620496 (Gamma3), AM493254 (Delta3), AF328857 (Delta1a/Delta1b), AF328856 (*Ca.* Thiosymbion); note that the Delta1a and Delta1b symbionts were represented by AF328857 as they share highly similar regions in their SSU sequences[2]). The results showed nearly identical community structures in all samples as compared to the SCG-based analyses (Supplementary Figure S4; cf. Figure 2). The only difference was that the relative abundances of the spirochete symbiont appeared slightly greater in the SSU-based estimation than in the SGC-based estimation. This minor difference is possibly due to the interspecific variation in the number of SSU among Bactria[3-5], which may be present among the symbionts of *O. algarvensis*.

#### 1.3. Symbiont 16S rRNA gene sequences indicated a linkage between haplotypes of *Candidatus* Thiosymbion and host mitochondria

As an initial assessment of partner fidelity, we assembled SSU sequences of symbionts in each *O. algarvensis* sample, and searched for SNPs that are characteristic to certain host COI-haplotypes and locations. Only the *Ca.* Thiosymbion symbiont showed a SNP site in the SSU sequences that was linked to the host COI-haplotype but not to locations (Supplementary Figure S9). No other symbionts showed SNPs in SSU sequences that could be linked to COI-haplotype or location.

#### 1.4. Reconstruction of mitochondria and symbiont phylogenies using a deterministic genotyping approach to SNP-identification

We identified SNPs in mitochondrial genome (mtDNA) and symbiont genomes using a deterministic genotyping approach to phylogenetic reconstruction, prior to the probabilistic approach to SNP-identification (see 'Identification of single nucleotide polymorphisms and phylogenetic reconstruction' in the Methods). For SNP-identification by deterministic genotyping, the same symbiont- and mitochondrial-reads were analyzed with the SNIPPY pipeline v3.2 (https://github.com/tseemann/snippy), with the same reference genomes of mtDNA and symbionts as described in the main article. Genotypes were first called at all reference nucleotide positions with a minimum coverage of 5× for each metagenome. Core SNP sites were subsequently identified among genotype-called sites that were covered ≥5× in all metagenomes. Similar to our procedures using the probabilistic approach, when no core SNP site was found, metagenomes with insufficient genotype data were excluded from phylogenetic analyses, using a cut-off of lateral coverage (i.e. % reference sites with coverage ≥5×; Supplementary Table S4). Phylogeny trees with bootstrap-support were computed from core SNP nucleotide alignment with IQ-TREE v1.5.5[6] using a general time reversible model with an ascertainment bias correction (GTR+ASC). Phylogenetic trees of the symbionts and mitochondria were visualized in the iTOL web tool[7].

Because the deterministic genotype-calling method (i) relies on deeply-sequenced genomic sites to account for sequencing errors, and (ii) identifies core SNP sites only at loci where all the metagenomes called genotypes, it performs conservative SNP-identification. However, it also imposes a limitation in the number of detectable SNP sites when low-coverage sequences are studied. In our study, this approach required us to exclude many metagenomes when no SNP site could be detected due to low sequence-coverages, or when certain symbionts were absent in the metagenome. Consequently, the number of metagenomes and SNPs we could include in downstream analyses were substantially less when using the deterministic genotyping approach, as compared to results using the probabilistic approach based on genotype probabilities (Supplementary Table S4). An exception was the spirochete symbiont, where slightly more SNPs were captured (99 sites) by the genotype-calling method than by the probabilistic approach (88 sites). This was likely due to exceptionally high genetic variability of the spirochete symbionts among host individuals (see Figure 4g), which resulted in many rare nucleotide variants that were removed by filtering of statistically insignificant SNP-sites by the probabilistic approach (SNP p-value > 0.01). Nevertheless, phylogenies based on limited SNPs and metagenomes using the deterministic genotyping method already showed overall clustering patterns comparable to the ones using the probabilistic approach as shown in Figure 4 (Supplementary Figure S5). Specifically, the core-SNP trees reproduced the distinctive divergence of mitochondria and *Ca*. Thiosymbion based on host COI-haplotypes (Supplementary Figure S5a and S5b), the location-based clustering of Delta4 (Supplementary Figure S5f), and the clustering of Gamma3 from one B-host from Cavoli within a clade of A-hosts from Cavoli (Supplementary Figure S5c; magnified panel). Similar to the results based on the probabilistic SNP-identification, phylogenetic relationships of mitochondria and *Ca*. Thiosymbion within the same mtDNA lineage were also not resolved using the deterministic genotyping method (Supplementary Figure S6c and S6d).

##### 1.5. Estimation of the effective population size of symbionts per *O. algarvensis* individual based on genome-wide SNP abundance

We estimated the effective population size ($N_e$) of each symbiont species per host individual based on genome-wide SNP abundance. $N_e$ is defined as a hypothetical number of individuals in an idealized population with a certain quantity of interest being equal to the one that the actual population shows. In our case, the quantity of interest is the nucleotide diversity of a symbiotic bacterial population, and the ideal population is one in which all genetic mutations are neutral. Calculation of $N_e$ for haploid species such as Bacteria and Archaea are challenging due to the potential ubiquity of genetic sites under selection and selective sweeps[8]. For simplicity, we followed one of the methods employed by Bobay and Ochman[8] that uses the Watterson's estimator θ to calculate $N_e$ (according to the equation; $\theta = 2 N_e \mu$, where μ is the mutation rate[9]). Based on our observations on a phylogenomic tree of *O. algarvensis* symbionts among other representative bacterial genomes (Supplementary Figure S1), we made the assumption that the mutation rates of all symbionts in *O. algarvensis* are similar. Specifically, we generated a phylogenomic tree of *O. algarvensis* symbiont genomes with reference bacterial genomes for which mutation rates are available[10]. The phylogeny was calculated based on 25 universally distributed single-copy genes in Bacteria and Archaea as implemented in GToTree v1.5.38[11]. The resulting tree showed that (**i**) published mutation rates for bacterial isolates previously studied are all in the similar order of magnitude around $10^{-10}$ mutation per site per generation, (**ii**) the mutation rates and branch lengths do not show a correlation within this range, and (**iii**) branch lengths to all symbionts of *O. algarvensis* are within the range of those to the bacterial isolates with published mutation rates. Consequently, we estimated the mutation rate μ of all symbionts as the average of the published mutation rates, $2.73\times10^{-10}$ per site per generation. For each symbiont in each of the 80 individuals, Watterson's θ was computed using the R package pegas v0.14[12] based on (i) the number of SNP-sites within an individual host and (ii) the number of sequences represented by the mean sequence coverage, as previously described[13]. The θ and $N_e$ were not calculated when mean sequence coverage of a symbiont in a given host individual was less than 5×, to ensure that each estimation of $N_e$. was based on sufficient data.

$N_e$ estimates of *O. algarvensis* symbionts were on the order of $10^4$ - $10^5$ (Supplementary Figure S8). $N_e$ estimates varied among the symbionts, but did not show any link to their partner fidelity levels indicated in this study (Table 1).

$N_e$ reflects the size of an actual population and fluctuations of genetic diversity over time. Specifically, in the case of endosymbionts, such as those associated with *O. algarvensis*, we can expect that the $N_e$ per host individual is determined by (**i**) the number of cells that colonise a new host generation, regardless of whether they are transferred via vertical transmission or horizontal transmission, (**ii**) the genetic diversity of the source population(s), (**iii**) internal bacterial growth after establishing a symbiosis, and/or (**iv**) the presence or absence of ongoing horizontal symbiont acquisition from an external source. Although it is not feasible to disentangle individual drivers of $N_e$ from simple comparisons of $N_e$ between symbionts, $N_e$ can be linked to the importance of certain drivers if other knowledge exists related to these drivers (see Supplementary text 1.6). For example, the relatively large $N_e$ estimates of *Ca*. Thiosymbion among the symbionts may stem from its numerical dominance in an adult host individual and in the inoculant during vertical transmission, as our study provides a strong support for vertical transmission of this symbiont (see discussion in the main article and Supplementary text 1.6). Other explanations are however possible for estimated $N_e$ values. The elucidation of drivers for $N_e$ will require the identification of the symbiont transmission mode, cell counts at the establishment of symbiosis and in an adult host, and genetic diversity of free-living symbionts if they are taken up by the host from the environment.

##### 1.6. Transmission modes of symbionts in *O. algarvensis*

The SNP-based phylogeny of *Ca.* Thiosymbion showed divergence into two clades linked to two mitochondrial lineages. This phylogenetic divergence was observed in both of the study locations. The strong host-symbiont fidelity observed for *Ca.* Thiosymbion can in theory arise through two alternative scenarios: (**i**) Coupling of A- and B-lineages between *O. algarvensis* mitochondria and the symbiont is mediated by genotype-dependent partner choice during horizontal symbiont acquisition, or (**ii**) stringent vertical symbiont transmission. For scenario (**i**), host mitochondrial lineages must have phenotypic differences that favour specific symbiont genotypes. Such phenotypic differentiation is likely maintained in the nuclear genome of the host populations, as the mitochondrial genomes (mtDNAs) of the two lineages share the high average nucleotide identity at 99.3%. Sexually reproducing populations homogenise genetic material among individuals via genetic recombination. Thus, establishing the linkage between host mtDNA and *Ca*. Thiosymbion lineages requires the co-occurring hosts with mitochondrial lineages A and B (A- and B-hosts) to be two reproductively-isolated populations. This scenario is unlikely because reproductive isolation at a mtDNA divergence of 0.7%, as we observed for *O. algarvensis*, is highly uncommon in clitallate annelids; e.g. Sympatric reproductive isolation in *Lumbricus rubellus* has only been observed with mtDNA sequence divergence of at least one order of magnitude higher[14,15].

The more likely scenario is stringent vertical transmission of the symbiont along with maternal mitochondria (**ii**); a mechanism that has been shown for many animal-bacterial symbioses[16-18]. The congruence of host- and symbiont-genealogies and a strong positive correlation of pairwise genetic distances between host mtDNA and *Ca*. Thiosymbion are all concordant with what is expected from stringent vertical transmission of this symbiont. Other previous genomic and morphological studies also corroborate vertical transmission of *Ca*. Thiosymbion: (i) Genetic and proteomic studies on *Ca*. Thiosymbion associated with *O. algarvensis* observed abundant transposases and their expression[19,20], one of the key signs of symbiont genome evolution upon the establishment of stringent vertical transmission[21,22]; and (ii) smearing of the primary *Ca.* Thiosymbion symbiont onto the egg of gutless annelids during ovipositioning was shown in microscopy observations[23-25]. Using the population-level phylogenomic approach for the first time, our study adds strong evidence for vertical transmission of *Ca*. Thiosymbion in *O. algarvensis.*

It is possible that symbionts other than *Ca.* Thiosymbion are similarly co-transmitted vertically, because vertical symbiont transmission often evolves as host-driven adaptation[26]. We speculate that the Gamma3 symbiont is inherited primarily by vertical transmission, as its phylogeny indicated (i) genetic differentiation partly explained by host mitochondrial divergence, and (ii) only a rare case of host switching between mitochondrial lineages in sympatric hosts. Vertical transmission may also be at work for the Delta1a and Delta1b symbionts, for which we observed notable patterns of their prevalence between A- and B-hosts (Figure 2). Estimates of the effective population sizes per host individual ($N_e$) for the Gamma3, Delta1a and Delta1b symbionts were lower than $N_e$ for *Ca*. Thiosymbion (Supplementary Figure S8; Supplementary text 1.5), which are in agreement with potential population-size bottlenecks associated with their vertical transmission. Previous studies have reported abundant transposases in genomes and proteomes of Gamma3 and Delta1a symbionts, suggesting some degree of host restriction in their lifestyle[19,20]. However, different degrees of mitochondria-symbiont genetic co-divergence among these symbionts suggest that they are inherited with varying relative importance of vertical transmission compared to horizontal transmission in the spectrum of mixed-mode transmission.

The location-based genetic divergence for the Delta4 symbiont suggests that this symbiont is horizontally transmitted within local host populations. There are possible hypotheses explaining the location-based patterns of the Delta4 symbiont: (**i**) Host individuals take up the Delta4 symbiont from free-living bacterial populations that are genetically differentiated between locations, (**ii**) co-occurring *O. algarvensis* individuals often exchange symbionts locally between A- and B-hosts (‘lineage switching’), or (**iii**) nuclear genomes of the hosts from the two locations are genetically well-differentiated and favours specific Delta4 genotypes during their horizontal acquirement from the environment.

For the first explanation (**i**), if Delta4 symbionts were horizontally acquired from a large environmental genetic pool of free-living bacterial populations, we could expect a larger genetic variation between host individuals for the Delta4 symbiont than that of *Ca*. Thiosymbion, because the *Ca*. Thiosymbion symbiont likely experiences population bottlenecks during vertical transmission. The Delta4 would be sampled anew from a diverse environmental population, which would lead to high inter-individual genetic variation. Interestingly, however, genetic variation of the Delta4 symbiont between host individuals was as low as that of *Ca*. Thiosymbion within the same host COI-haplotype and location (Figure 4a and 4f "Within" terms). In general, genetic diversity of free-living bacterial species in the coastal environment can be high; e.g., 16S ribosomal RNA gene (SSU) sequences of the sulphate-reducing *Desulfobacterium* genus show high conspecific diversity within a coastal saltmarsh sediment sample[27]. As no comparable SNP data was available for free-living Delta4 populations at our study sites, we tested for signals of nucleotide diversity in the SSU sequences of the Delta4 symbiont, but could not detect any (Supplementary Figure S9). It is unlikely that large and diverse free-living Delta4 populations, if they existed, could maintain the low genetic diversity of the host-associated Delta4 symbionts among sympatric host individuals. Therefore it is unlikely that the genetic differentiation of free-living Delta4 symbionts between locations can explain our observation.

The second explanation (**ii**) i.e. frequent local lineage-switching of the Delta4 symbiont is more likely. The low $N_e$ estimates for the Delta4 symbiont indicate that the low genetic diversity of the symbiont is not only between host individuals but also within the host individuals (Supplementary Figure S8). Frequent local lineage-switching of symbionts, while maintaining their low genetic diversity within and between host individuals, may be mediated by biparental or paternal symbiont transmission that are observed in other symbioses (e.g. biparental transmission of nephridial symbionts in the earthworm *Aporrectodea tuberculata*[28], and male-borne transmission of symbionts in aphids and tsetse fly[29,30]). Biparental and paternal symbiont transmission can facilitate regular symbiont exchanges between host lineages within a local host population, and maintain the low genetic diversity within host individual. The third explanation (**iii**) is also possible, but requires a study to identify host nuclear genome structuring between locations, as well as identifying genomic loci in host and Delta4 that are responsible for a potential partner choice.

The spirochete symbiont showed neither phylogenetic patterns linked to host mtDNA lineage, nor genetic co-divergence with host mtDNA. Such a lack of phylogenetic evidence for partner fidelity indicates that the spirochete is regularly acquired from the environment with horizontal transmission[31-33]. Genetic variations of the spirochete symbiont between sympatric individuals of A- or B-hosts was more than 2-fold greater than that of all other symbionts (Figure 4g "Within"), and $N_e$ estimates for the spirochete symbiont were the highest among *O. algarvensis* symbionts (Supplementary Figure S8). Considering that the spirochete symbiont did not show a location effect on the phylogenetic clustering patterns (Figure 4g), our results suggest that the spirochete symbionts derives from genetic pools that are so large and diverse that they are undifferentiated between the two locations. Such genetic pools are likely free-living populations in the environment. However, our phylogenetic result cannot rule out occasional vertical transmission of the spirochete symbiont between host generations, because symbiont phylogenies will not show evidence for vertical transmission events if horizontal transmission is sufficiently frequent or regular mode of acquisition.

18 Allen, J. M., Reed, D. L., Perotti, M. A. & Braig, H. R. Evolutionary Relationships of "*Candidatus* Riesia spp.," Endosymbiotic *Enterobacteriaceae* Living within Hematophagous

Primate Lice. *Applied and Environmental Microbiology* **73**, 1659-1664, doi:10.1128/aem.01877-06 (2007).
19 Woyke, T. *et al.* Symbiosis insights through metagenomic analysis of a microbial consortium. *Nature* **443**, 950-955, doi:10.1038/nature05192 (2006).
20 Kleiner, M., Young, J. C., Shah, M., VerBerkmoes, N. C. & Dubilier, N. Metaproteomics Reveals Abundant Transposase Expression in Mutualistic Endosymbionts. *mBio* **4**, e00223-00213, doi:10.1128/mBio.00223-13 (2013).
21 Moran, N. A. Accelerated evolution and Muller's rachet in endosymbiotic bacteria. *Proceedings of the National Academy of Sciences of the United States of America* **93**, 2873-2878, doi: 10.1073/pnas.93.7.2873 (1996).
22 Moran, N. A. & Plague, G. R. Genomic changes following host restriction in bacteria. *Current Opinion in Genetics & Development* **14**, 627-633, doi:10.1016/j.gde.2004.09.003 (2004).
23 Giere, O. Ecology and Biology of Marine Oligochaeta – an Inventory Rather than another Review. *Hydrobiologia* **564**, 103-116, doi:10.1007/s10750-005-1712-1 (2006).
24 Giere, O. & Langheld, C. Structural organisation, transfer and biological fate of endosymbiotic bacteria in gutless oligochaetes. *Marine Biology* **93**, 641-650, doi:10.1007/BF00392801 (1987).
25 Schimak, M.-P. *Transmission of bacterial symbionts in the gutless oligochaete Olavius algarvensis*, University of Bremen, (2016).
26 Sachs, J. L., Skophammer, R. G. & Regus, J. U. Evolutionary transitions in bacterial symbiosis. *Proceedings of the National Academy of Sciences* **108**, 10800-10807, doi:10.1073/pnas.1100304108 (2011).
27 Klepac-Ceraj, V. *et al.* High overall diversity and dominance of microdiverse relationships in salt marsh sulphate-reducing bacteria. *Environmental Microbiology* **6**, 686-698, doi:10.1111/j.1462-2920.2004.00600.x (2004).
28 Paz, L.-C., Schramm, A. & Lund, M. B. Biparental transmission of *Verminephrobacter* symbionts in the earthworm *Aporrectodea tuberculata* (Lumbricidae). *FEMS Microbiology Ecology* **93**, fix025, doi:10.1093/femsec/fix025 (2017).
29 Moran, N. A. & Dunbar, H. E. Sexual acquisition of beneficial symbionts in aphids. *Proceedings of the National Academy of Sciences* **103**, 12803-12806, doi:10.1073/pnas.0605772103 (2006).
30 De Vooght, L., Caljon, G., Van Hees, J. & Van Den Abbeele, J. Paternal Transmission of a Secondary Symbiont during Mating in the Viviparous Tsetse Fly. *Molecular Biology and Evolution* **32**, 1977-1980, doi:10.1093/molbev/msv077 (2015).
31 Douglas, A. E. & Werren, J. H. Holes in the Hologenome: Why Host-Microbe Symbioses Are Not Holobionts. *mBio* **7**, e02099-02015, doi:10.1128/mBio.02099-15 (2016).
32 Bright, M. & Bulgheresi, S. A complex journey: transmission of microbial symbionts. *Nature Reviews Microbiology* **8**, 218-230, doi:10.1038/nrmicro2262 (2010).
33 Won, Y.-J., Jones, W. J. & Vrijenhoek, R. C. Absence of Cospeciation Between Deep-Sea Mytilids and Their Thiotrophic Endosymbionts. *Journal of Shellfish Research* **27**, 129-138, doi:10.2983/0730-8000(2008)27[129:AOCBDM]2.0.CO;2 (2008).

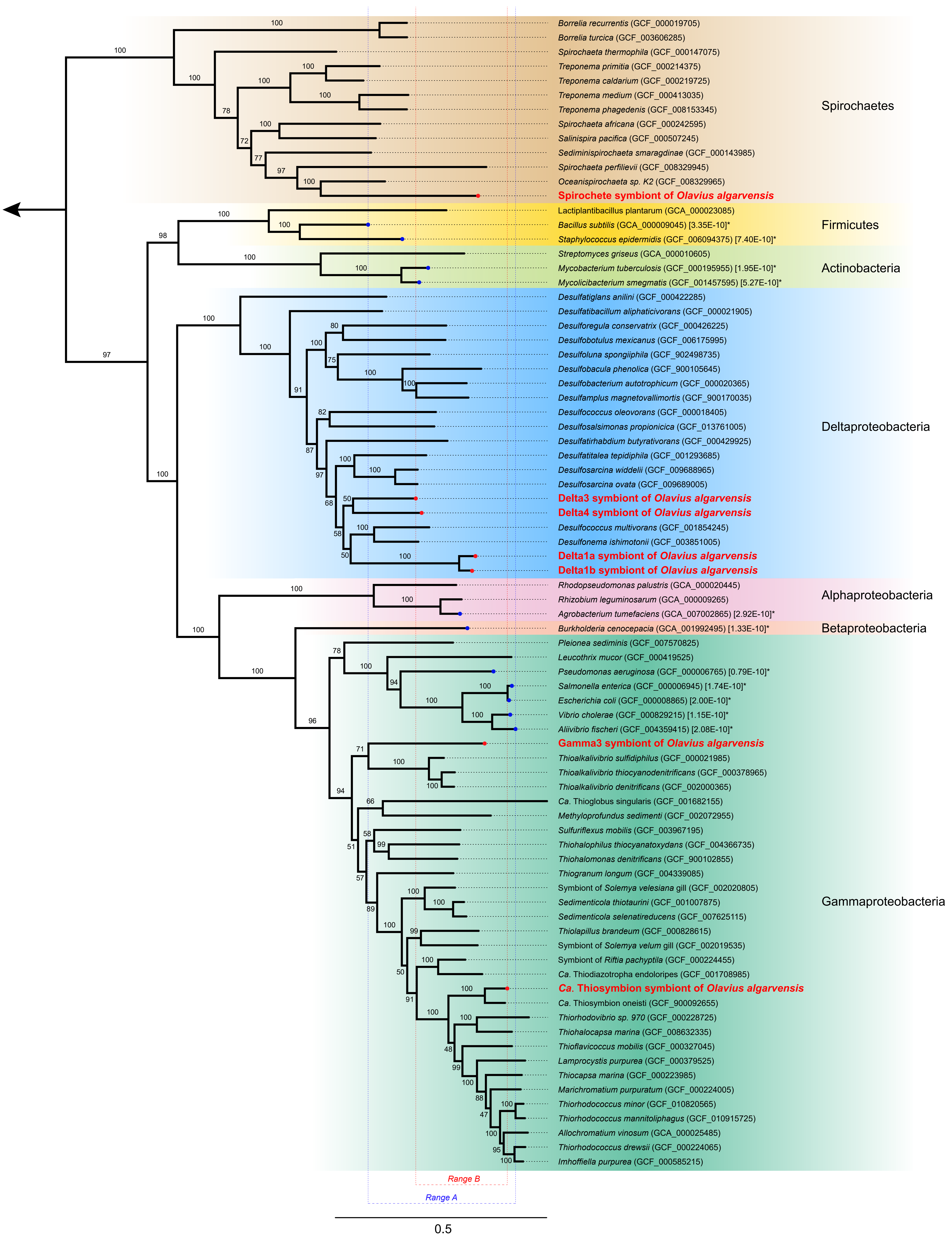


**Supplementary Figure S1** Phylogenomic tree of symbionts in *Olavius algarvensis* in relation to reference bacterial genomes. The phylogeny is calculated based on amino acid sequences of 25 protein coding genes that are universally present in Bacteria and Archaea. NCBI Assembly accession numbers are indicated in brackets. The outgroup is the archaeon *Nitrososphaera viennensis* (GCA_000698785), indicated with the arrow. The scale bar indicates 0.5 amino acid replacement per amino acid site. The genomes of *O. algarvensis* symbionts are highlighted with red bold fonts and red dots at the branch terminal. Asterisks and blue dots denote reference bacteria for which mutation rates, as indicated in square brackets [mutation per nucleotide site per generation], are available (Lynch et al. 2016). Note that the range of branch lengths for these reference bacteria (Range A) encompasses the range of branch lengths for the *O. algarvensis* symbiont genomes (Range B). Branch support values indicate IQ-TREE Ultrafast Bootstrap estimates with 1,000 times replication.

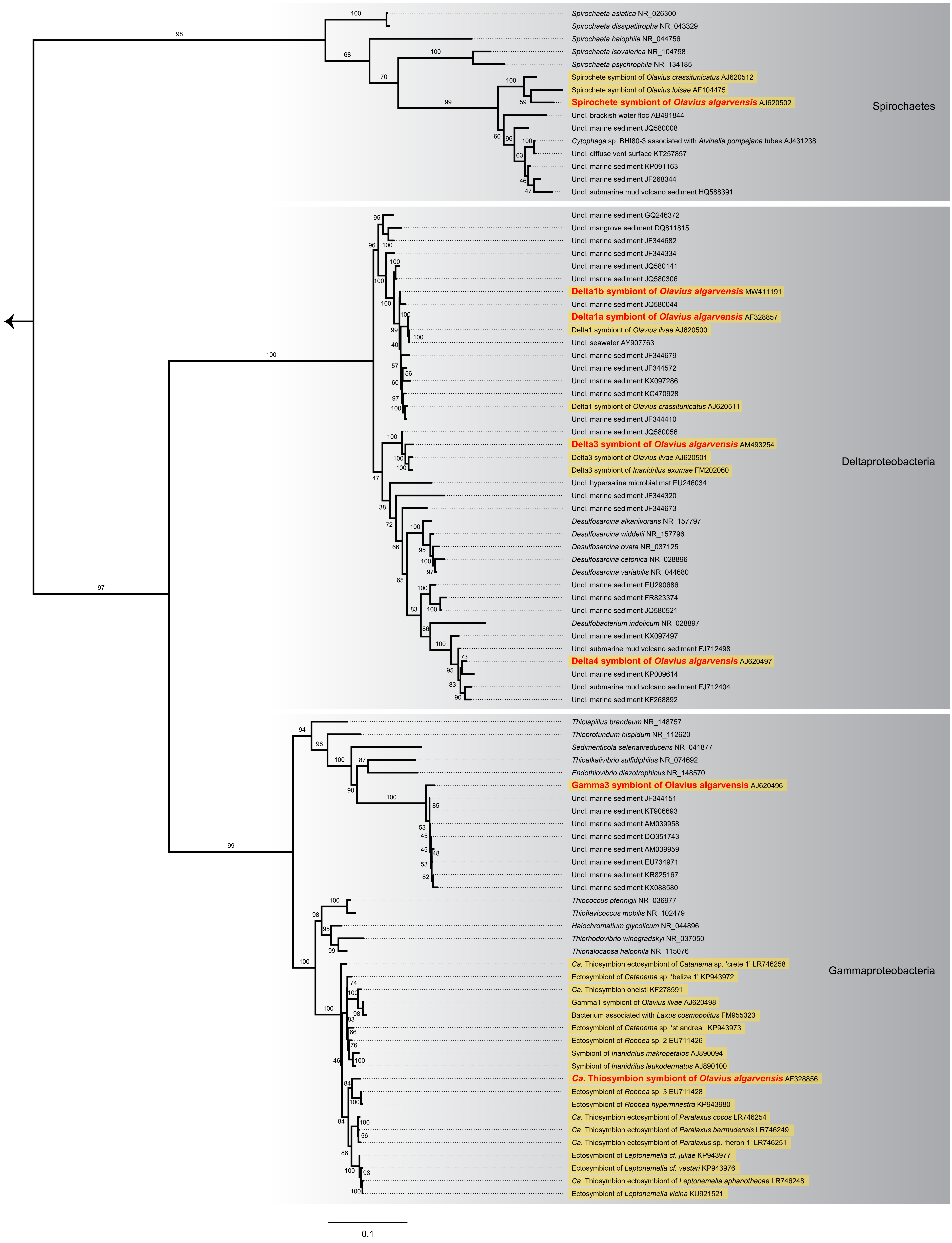


**Supplementary Figure S2** Phylogeny of 16S rRNA gene sequences for symbionts in *Olavius algarvensis* in relation to reference bacterial sequences. NCBI sequence accession numbers are indicated in the labels. The outgroup is the archaeon *Nitrososphaera viennensis* (FR773157), indicated with the arrow. The scale bar indicates 0.1 base pair replacement per nucleotide. The sequences of *O. algarvensis* symbionts are highlighted with red bold fonts. Sequences of bacteria associated with host animals are highlighted with yellow labels. "Uncl." denotes an uncultured bacterium. Branch support values indicate IQ-TREE Ultrafast Bootstrap estimates with 1,000 replications.

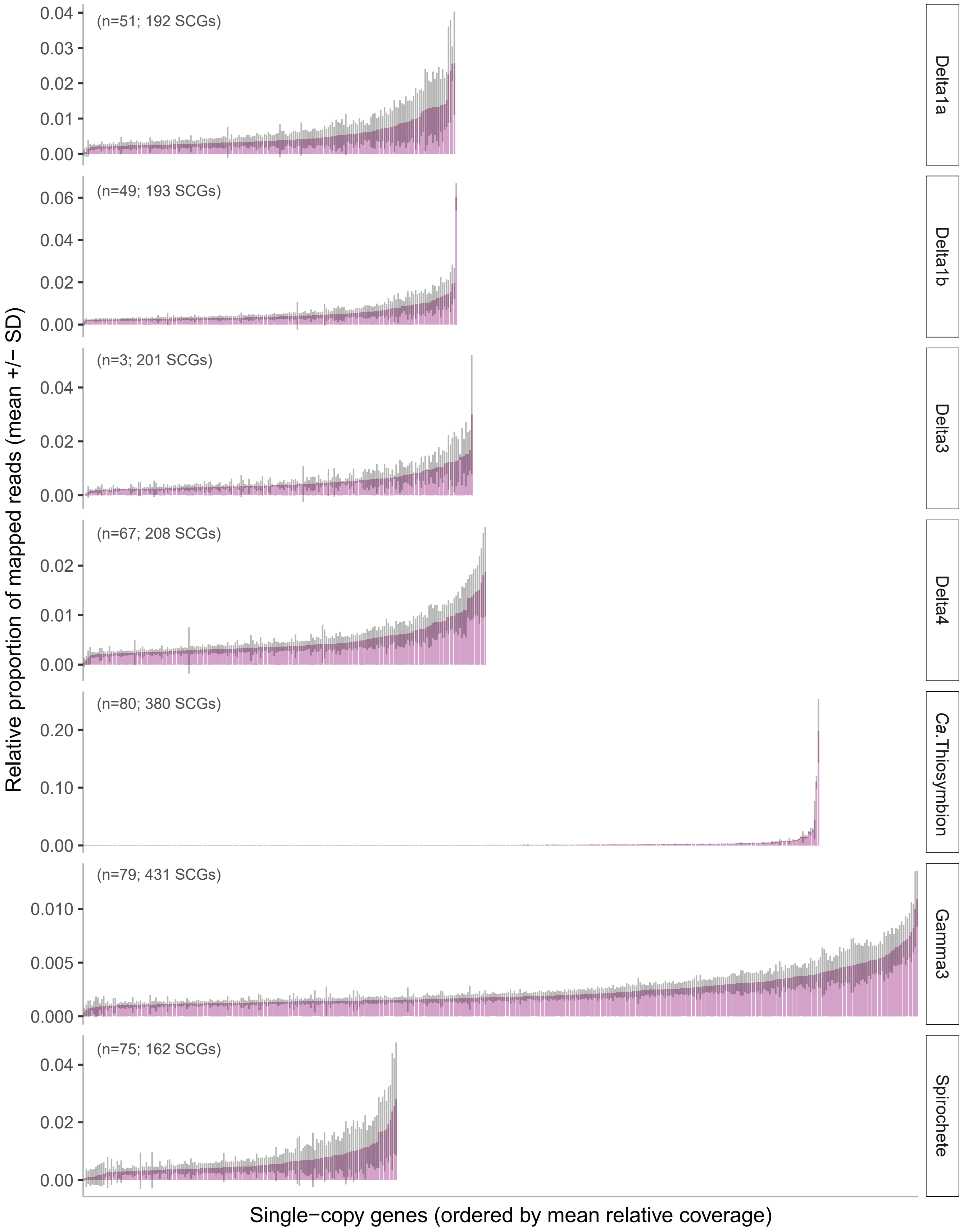


**Supplementary Figure S3** Relative proportions of mapped reads to each of the single-copy genes (SCGs) identified a small subset of SCGs with erroneously high read coverages. Plots are shown as the mean proportion of read coverage (pink bars) ± standard deviation (SD; grey bars) per gene. The number of replicate samples and total number of reference SCG sequences included in reference are shown in brackets.

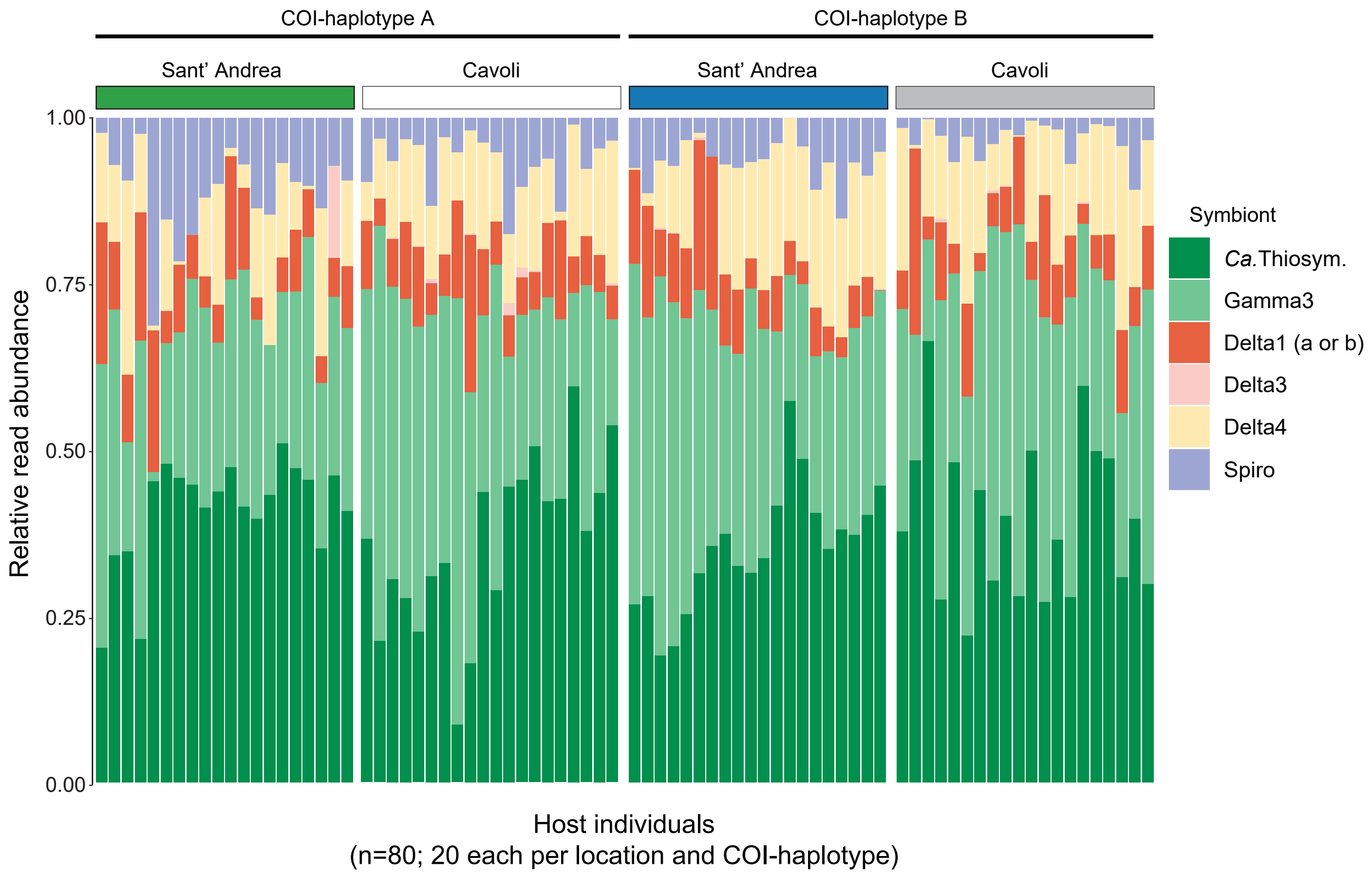


**Supplementary Figure S4** Symbiont composition of individual *O. algarvensis* samples of two COI-haplotypes (A and B) from two locations (Sant' Andrea and Cavoli). Relative abundance was estimated based on 16S rRNA gene sequences mapped to a collection of reference sequences derived from *O. algarvensis* symbionts. Note that Delta1a and Delta1b symbionts are pooled and represented as "Delta1" because they share identical regions of 16S rRNA genes and cannot be distinguished by read-mapping. "*Ca.*Thiosym." refers to *Candidatus* Thiosymbion.

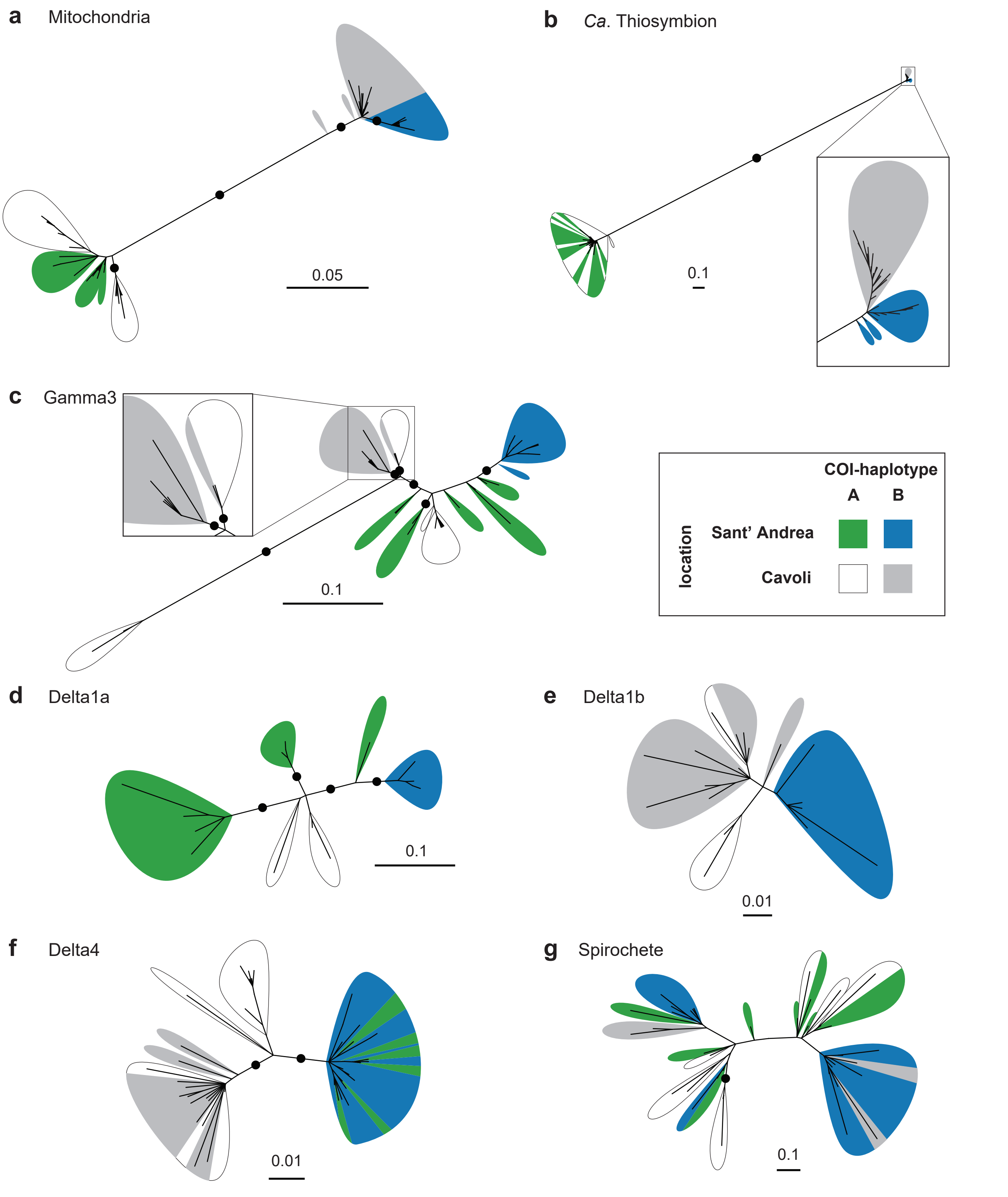


**Supplementary Figure S5** Core-SNP phylogenies using the deterministic approach to genotype-calling (SNIPPY v3.2) for **a**; mitochondria (121 SNPs, 76 samples), **b**; *Ca*. Thiosymbion (391 SNPs, 76 samples), **c**; Gamma3 (73 SNPs, 72 samples), **d**; Delta1a (72 SNPs, 16 samples), **e**; Delta1b (57 SNPs, 25 samples), **f**; Delta4 (177 SNPs, 45 samples), and **g**; Spirochete (99 SNPs, 40 samples). Scale bars indicate substitution per SNP-site. Bootstrap support values >95% are shown in black circles. Supports for branches internal to each coloured leaf are omitted for visibility. Note: In the magnified panel in (**c**), the Gamma3 symbionts in five A-hosts from Cavoli (white) shared the identical core SNPs and thus are represented by a single node in the graph.

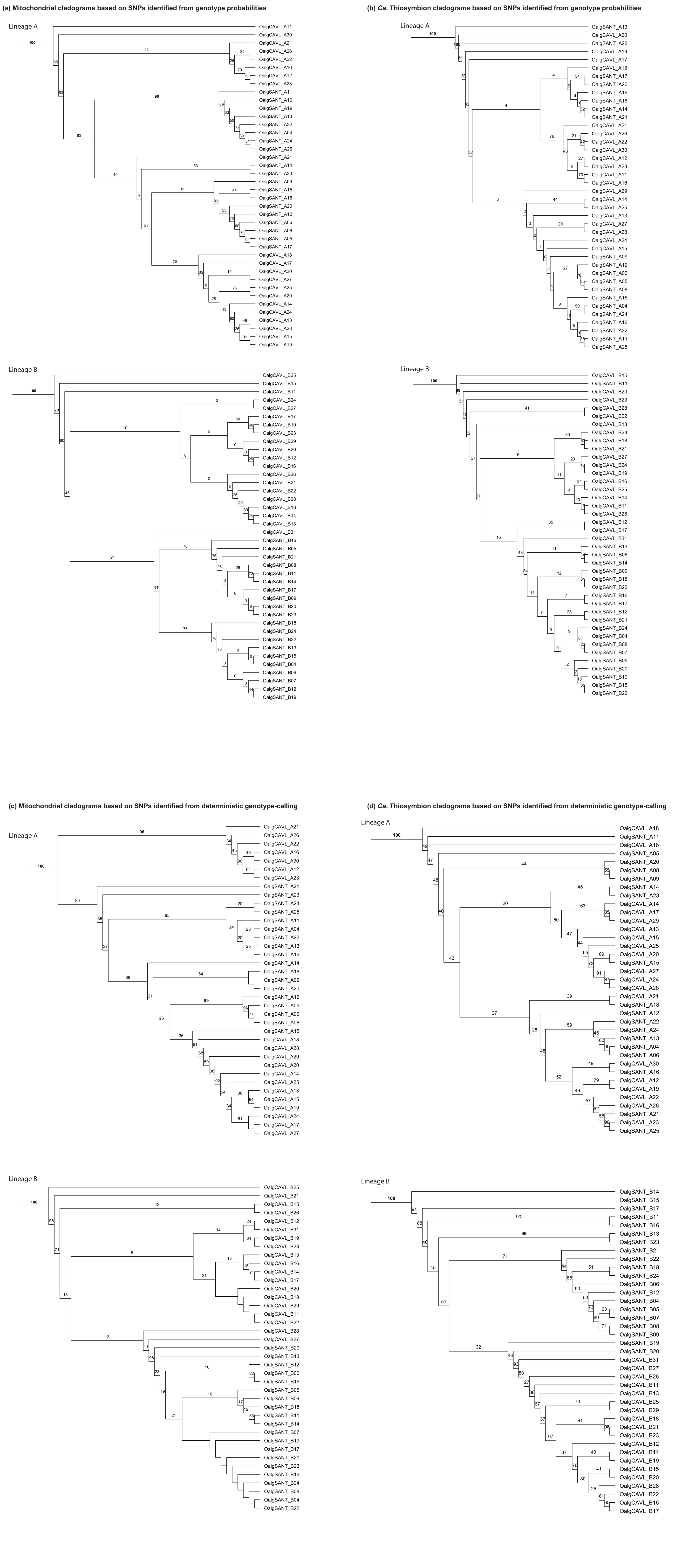


**Supplementary Figure S6** Sub-clades within the major mitochondrial lineages A and B were poorly supported for host mitochondrial phylogenies using a probabilistic approach to SNP identification (**a**) and deterministic genotype-calling (**c**), as well as for *Ca*. Thiosymbion phylogenies using the probabilistic approaching to SNP identification (**b**) and deterministic genotype calling (**d**). Bootstrap support values above 95% are highlighted with bold fonts. Branches without clade supports are due to polychotomy. Uniform total branch lengths are transformed proportionally to branch-lengths in the phylogeny shown in Figure 3 and Supplementary Figure S5 for improving the visibility of bootstrap support values.

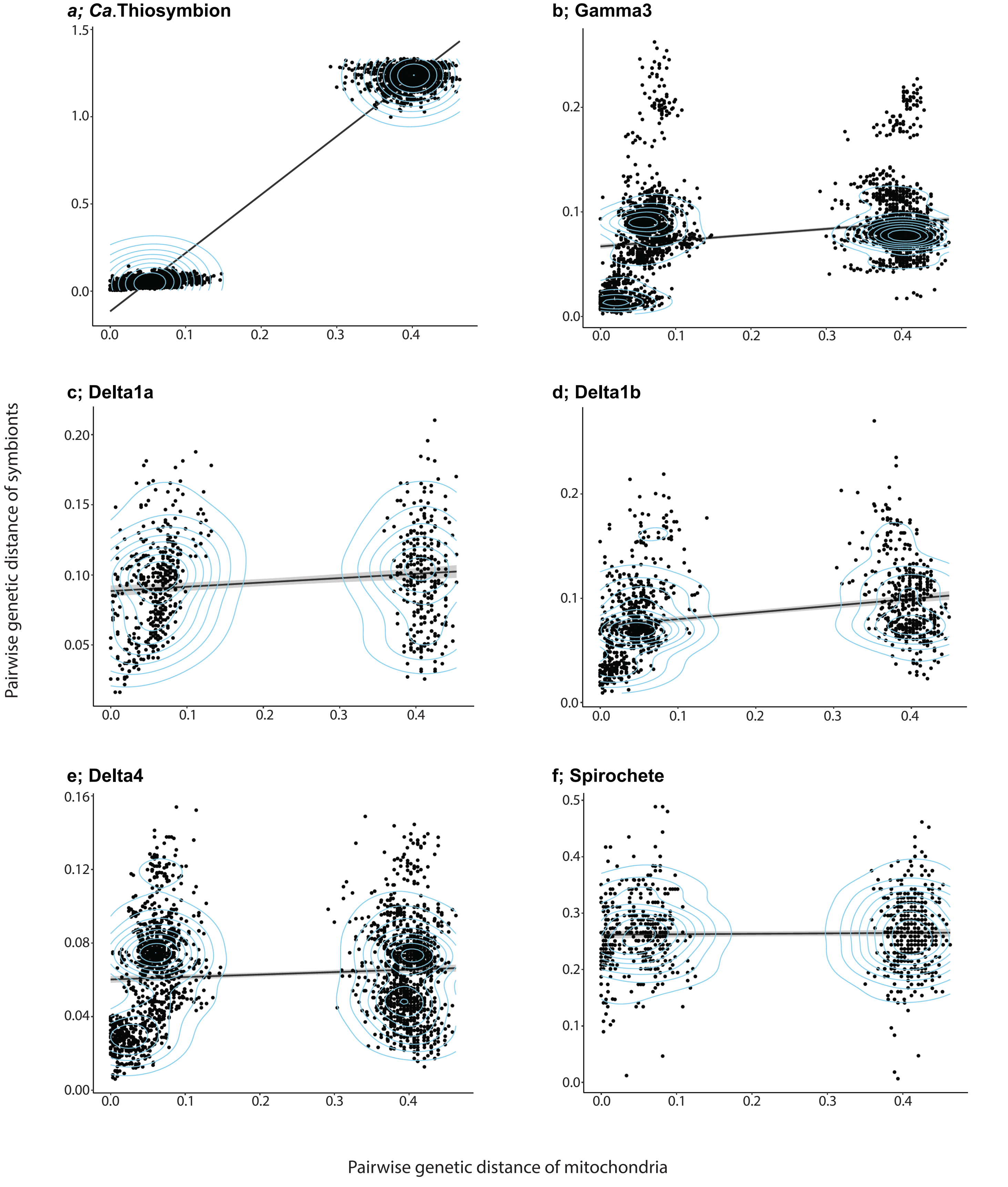


**Supplementary Figure S7** Correlation between pairwise genetic distance of mitochondria (x-axis) and pairwise genetic distance of each symbiont (y-axis). Each dot represents a set of two individuals. Genetic distances were measured using NGSdist based on posterior genotype-probabilities calculated with ANGSD. Linear regression lines and 95% confidence intervals are shown with a black line and grey ribbon, respectively. Note the different regression patterns, with *Candidatus* Thiosymbion showing the strongest positive correlation, and other symbionts showing small or no positive correlations between the mitochondrial distance and symbiont distance. Plot densities are indicated with pale blue contours, and Gamma3 and Delta4 respectively displayed three and four centres of plot density, indicating the presence of factor(s) other than mitochondrial divergence that explain the genetic divergence of these two symbionts.

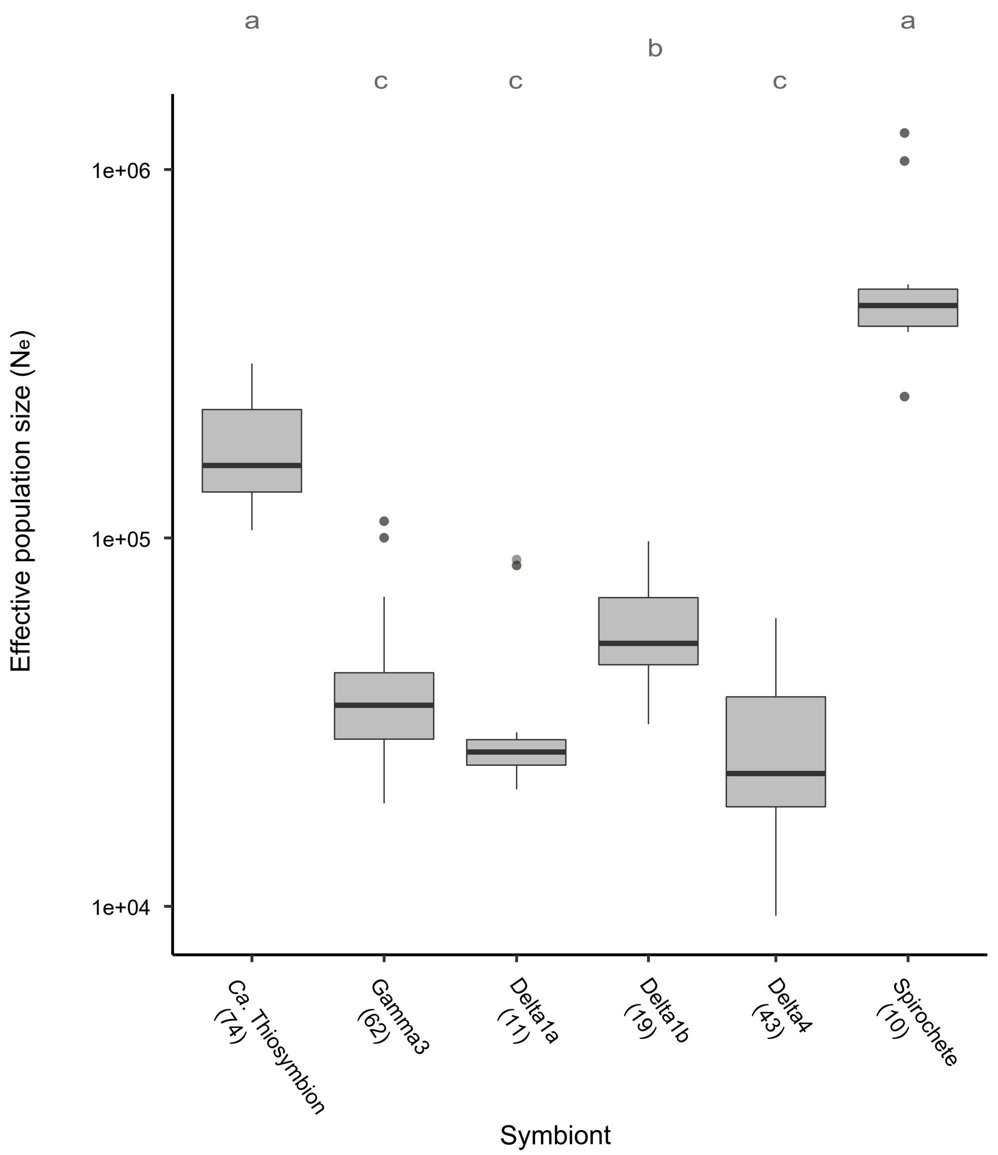


**Supplementary Figure S8** Effective population size (Ne) estimates of the symbiont per *Olavius algarvensis* individual. Estimation was based on the number of genome-wide segregation sites (SNPs), the average sequence coverage and an assumed mutation rate (see Supplementary text 1.5). Numbers in brackets indicate numbers of replicates for each symbiont. Note that the y-axis is in log-scale. Thick horizontal lines and grey boxes respectively indicate the median and interquartile range (IQR) of observations. Vertical lines show the IQR ± 1.5 IQR range, and outliers out of this range are shown as circles. Letters on the top indicate a summary of pairwise statistical comparisons based on a p-value cutoff of 0.05 (post-hoc Dunn multiple comparison tests with the Benjamini-Hochberg p-value adjustment. Prior Kruskal-Wallis rank sum test indicated p-value = 2.2e-16, $\chi^2$ = 172.89, df = 5).

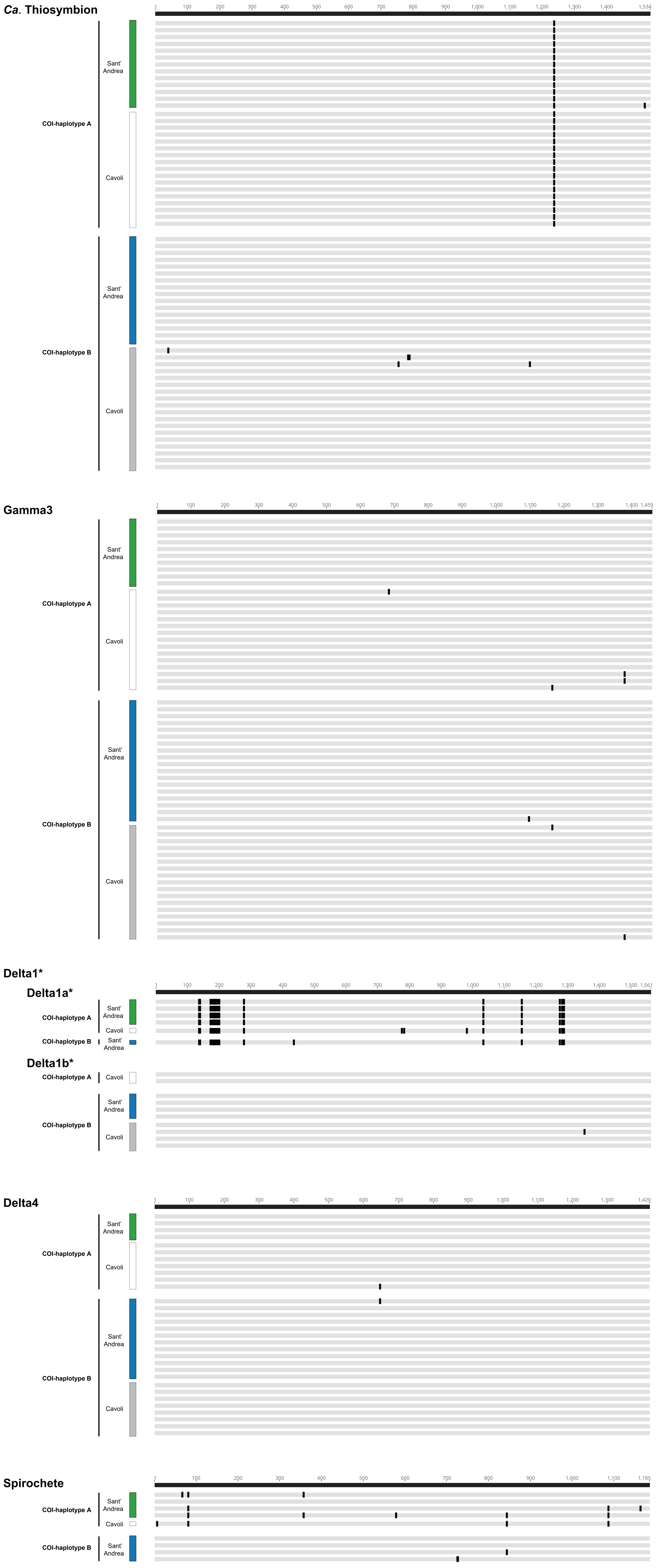


**Supplementary Figure S9** Sequence alignments of 16S rRNA genes of *O. algarvensis* symbionts (from top to bottom; *Candidatus* Thiosymbion, Gamma3, Delta1 (Delta1a and Delta1b), Delta4 and spirochetes). Within each symbiont species, black lines denote single nucleotide polymorphisms among non-variable sites indicated by grey boxes. COI-haplotypes (A and B) and sampling locations (Sant' Andrea and Cavoli) of the host *O. algarvensis* worms are indicated with coloured boxes on the left. Among the symbiont species, 16S rRNA gene sequences were successfully assembled from many host individuals for *Ca*. Thiosymbion, Gamma3 and Delta4, while assembly of 16S rRNA gene sequences for the Delta1a, Delta1b and spirochete symbionts was unsuccessful in many metagenomes, due to their low relative abundances (or absence) and low sequence-coverages. Note that only the *Ca.* Thiosymbion symbiont show a distinctive SNP site that clearly separates COI-haplotypes A and B of the host, but no other symbionts show SNPs that can be linked to host COI-haplotype or location.

*Delta1 sequences were classified into Delta1a and Delta1b symbionts by cross-checking with the presence/absence of Delta1a and Delta1b symbionts in each host assessed by quantification of species-specific single-copy gene sequences (Figure 2 in the main article). Note Delta1a and Delta1b sequences share long non-variable regions.

**Supplementary Table S1** Summary statistics of the reference metagenome-assembled genomes of *O. algarvensis* symbionts

| **Reference genome** | ***Ca.* Thiosymbion** | **Gamma3** | **Delta1a** | **Delta1b** | **Delta3** | **Delta4** | **Spirochete** |
|---|---|---|---|---|---|---|---|
| Specimen ID | OalgB6SA | OalgB6SA | OalgA4SA | OalgB6SA | OalgB2SA | OalgB6SA | OalgB6SA |
| NCBI accession number | GCA_905176695 | GCA_905176675 | GCA_902749705 | GCA_902749685 | GCA_903231395 | GCA_905176665 | GCA_905176685 |
| COI-haplotype | B | B | A | B | B | B | B |
| Collection site | Sant’ Andrea | Sant’ Andrea | Sant’ Andrea | Sant’ Andrea | Sant’ Andrea | Sant’ Andrea | Sant’ Andrea |
| Genome size (bp) | 3,819,172 | 4,272,394 | 10,617,271 | 9,888,100 | 5,583,787 | 5,465,468 | 2,393,688 |
| #contigs | 8,736 | 1,048 | 4,867 | 1,617 | 401 | 286 | 515 |
| #contigs >=1kbp | 927 | 268 | 1,427 | 523 | 131 | 98 | 135 |
| #contigs >=5kbp | 68 | 179 | 705 | 308 | 109 | 71 | 68 |
| #contigs >=10kbp | 2 | 143 | 311 | 227 | 92 | 61 | 52 |
| #contigs >=50kbp | 0 | 10 | 1 | 49 | 38 | 38 | 13 |
| Largest contig | 11,494 | 166,547 | 64,030 | 206,304 | 424,721 | 333,300 | 174,819 |
| N50 | 1,978 | 23,651 | 10,152 | 44,270 | 76,958 | 121,536 | 45,935 |
| N75 | 344 | 13,833 | 5,349 | 20,618 | 46,596 | 64,281 | 15,534 |
| L50 | 551 | 50 | 307 | 66 | 21 | 13 | 15 |
| L75 | 1,616 | 108 | 668 | 148 | 44 | 28 | 36 |
| GC (%) | 55.99 | 55.62 | 49.45 | 48.10 | 54.22 | 53.99 | 47.40 |
| Completeness (%) | 82.20 | 98.24 | 95.81 | 96.13 | 92.90 | 99.35 | 97.60 |
| Contamination (%) | 0.43 | 3.14 | 3.55 | 2.58 | 0.00 | 1.29 | 8.00 |
| Strain heterogeneity (%) | 0 | 0 | 16.7 | 0 | 0 | 0 | 4.35 |
| Genome coverage (×) | 80 | 185 | 91 | 29 | 36 | 36 | 16 |

*N50/75 is the length (bp) for which the collection of all contigs of that length or longer covers at least 50/75% of the genome.

L50/75 is the minimal number of contigs that cover 50/75% of the genome.

**Supplementary Table S2** Symbionts in *O. algarvensis* have extremely low strain diversity within a host individual: (**a**) statistics of metagenomes used for the analysis with read-coverage of each symbiont genome, and (**b**) SNP-site density per 1kbp of each symbiont.

**a**

| ENA sequence accession (Specimen ID) | COI-haplotype | Location | Total data (Gb) | Coverage | | | | | | |
|---|---|---|---|---|---|---|---|---|---|---|
| | | | | Delta1a | Delta1b | Delta3 | Delta4 | *Ca.* Thiosymbion | Gamma3 | Spirochete |
| ERR3773751 (OalgA4SA) | A | Sant' Andrea | 5.7 | 91.1 | 0.0 | 0.0 | 56.2 | 62.0 | 115.6 | 5.5 |
| SRR5248183 (OalgA2SA) | A | Sant' Andrea | 11.8 | 14.4 | 0.0 | 0.0 | 24.8 | 72.4 | 35.9 | 7.7 |
| SRR5251647 (OalgA1SA) | A | Sant' Andrea | 17.2 | 17.1 | 0.0 | 0.0 | 22.4 | 44.6 | 32.4 | 14.3 |
| SRR6213993 (OalgA3SA) | A | Sant' Andrea | 25.4 | 41.7 | 0.0 | 0.0 | 74.0 | 157.8 | 119.0 | 34.3 |
| SRR5421031 (OalgA1CA) | A | Cavoli | 11.2 | 12.2 | 19.8 | 55.5 | 0.0 | 84.0 | 50.9 | 9.2 |
| SRR5421034 (OalgA2CA) | A | Cavoli | 25.9 | 31.7 | 0.0 | 0.0 | 79.2 | 161.4 | 151.7 | 25.5 |
| SRR5421607 (OalgA3CA) | A | Cavoli | 11.7 | 20.2 | 0.0 | 0.0 | 52.8 | 108.6 | 68.0 | 14.7 |
| ERR3773750 (OalgB6SA) | B | Sant' Andrea | 6.8 | 0.0 | 28.9 | 0.0 | 35.6 | 80.0 | 185.0 | 15.8 |
| SRR5248184 (OalgB1SA) | B | Sant' Andrea | 13.0 | 0.0 | 0.0 | 0.0 | 46.1 | 73.4 | 59.2 | 10.3 |
| SRR5248185 (OalgB2SA) | B | Sant' Andrea | 16.5 | 35.1 | 35.1 | 35.6 | 0.0 | 89.2 | 86.6 | 14.9 |
| SRR5248194 (OalgB3SA) | B | Sant' Andrea | 13.8 | 0.0 | 14.0 | 0.0 | 23.8 | 56.2 | 52.1 | 6.5 |

**b**

| ENA sequence accession (Specimen ID) | COI-haplotype | location | SNP-density (SNP/kbp) | | | | | | |
|---|---|---|---|---|---|---|---|---|---|
| | | | Delta1a | Delta1b | Delta3 | Delta4 | *Ca.* Thiosymbion | Gamma3 | Spirochete |
| ERR3773751 (OalgA4SA) | A | Sant' Andrea | 0.04 | n.a. | n.a. | 0.03 | 0.46 | 0.03 | 0.50 |
| SRR5248183 (OalgA2SA) | A | Sant' Andrea | 0.09 | n.a. | n.a. | 0.02 | 0.50 | 0.05 | 0.31 |
| SRR5251647 (OalgA1SA) | A | Sant' Andrea | 0.07 | n.a. | n.a. | 0.02 | 0.41 | 0.05 | 0.40 |
| SRR6213993 (OalgA3SA) | A | Sant' Andrea | 0.12 | n.a. | n.a. | 0.04 | 0.59 | 0.15 | 0.58 |
| SRR5421031 (OalgA1CA) | A | Cavoli | 0.19 | 0.15 | 0.03 | n.a. | 0.73 | 0.08 | 0.48 |
| SRR5421034 (OalgA2CA) | A | Cavoli | 0.10 | n.a. | n.a. | 0.03 | 0.55 | 0.13 | 1.00 |
| SRR5421607 (OalgA3CA) | A | Cavoli | 0.08 | n.a. | n.a. | 0.03 | 0.55 | 0.16 | 0.55 |
| ERR3773750 (OalgB6SA) | B | Sant' Andrea | n.a. | 0.06 | n.a. | 0.01 | 0.29 | 0.07 | 0.45 |
| SRR5248184 (OalgB1SA) | B | Sant' Andrea | n.a. | n.a. | n.a. | 0.03 | 0.46 | 0.08 | 0.41 |
| SRR5248185 (OalgB2SA) | B | Sant' Andrea | 0.09 | 0.09 | 0.02 | n.a. | 0.35 | 0.08 | 0.38 |
| SRR5248194 (OalgB3SA) | B | Sant' Andrea | n.a. | 0.09 | n.a. | 0.02 | 0.46 | 0.07 | 0.13 |
| ***average*** | | | ***0.10*** | ***0.10*** | ***0.02*** | ***0.03*** | ***0.49*** | ***0.09*** | ***0.47*** |

**Supplementary Table S3** Ratios between summed relative abundances of the gammaproteobacterial and deltaproteobacterial symbionts were consistent regardless of the locations (**a**), A- or B-hosts (**b**), or their combinations (**c**).

**a;** Comparison of Gammaproteobacteria/Deltaproteobacteria ratios between locations

| location | mean | SD | min. | median | max. | n |
|---|---|---|---|---|---|---|
| Cavoli | 3.02 | 1.11 | 1.21 | 2.68 | 5.60 | 40 |
| Sant' Andrea | 2.81 | 1.25 | 1.24 | 2.50 | 7.71 | 40 |

| | $\chi 2$ | df | p-value |
|---|---|---|---|
| Kruskal-Wallis test | 1.31 | 1 | 0.25 |

**b;** Comparison of Gammaproteobacteria/Deltaproteobacteria ratios between A- and B-hosts

| host type | mean | SD | min. | median | max. | n |
|---|---|---|---|---|---|---|
| A | 3.12 | 1.34 | 1.24 | 3.02 | 7.71 | 40 |
| B | 2.71 | 0.97 | 1.21 | 2.43 | 5.51 | 40 |

| | $\chi 2$ | df | p-value |
|---|---|---|---|
| Kruskal-Wallis test | 2.43 | 1 | 0.12 |

**c;** Comparison of Gammaproteobacteria/Deltaproteobacteria ratios among combinations of host types and location

| host type, location | mean | SD | min. | median | max. | n |
|---|---|---|---|---|---|---|
| A, Sant' Andrea | 3.19 | 1.57 | 1.24 | 3.17 | 7.71 | 20 |
| A, Cavoli | 3.06 | 1.09 | 1.38 | 2.70 | 5.60 | 20 |
| B, Sant' Andrea | 2.43 | 0.67 | 1.72 | 2.27 | 4.12 | 20 |
| B, Cavoli | 2.99 | 1.15 | 1.21 | 2.65 | 5.51 | 20 |

| | $\chi 2$ | df | p-value |
|---|---|---|---|
| Kruskal-Wallis test | 5.19 | 3 | 0.16 |

(SD; Standard deviation)

**Supplementary Table S4** A probabilistic approach to SNP-identification based on posterior genotype probabilities enabled analysis of increased numbers of host individuals (**a**) and based on more SNP-sites (**b**), compared to those using the deterministic approach based on called genotypes.

**a**; Number of host individuals

| Subject | Deterministic approach | Probabilistic approach |
|---|---|---|
| Mitochondria | 76 | 80 |
| *Ca*. Thiosymbion | 76 | 80 |
| Gamma3 | 72 | 80 |
| Delta1a | 16 | 37 |
| Delta1b | 25 | 46 |
| Delta4 | 45 | 67 |
| Spirochete | 40 | 41 |

**b**; Number of SNP sites and SNP density

| Subject | Genome size (bp) | Deterministic approach* | | | Probabilistic approach** | | |
|---|---|---|---|---|---|---|---|
| | | SNP sites | SNP density (SNP/kbp) | Cutoff | SNP sites | SNP density (SNP/kbp) | Cutoff |
| Mitochondria | 15,715 | 121 | 7.700 | (>80%) | 166 | 10.563 | |
| *Ca*. Thiosymbion | 3,819,172 | 391 | 0.102 | (>29%) | 2872 | 0.752 | |
| Gamma3 | 4,272,394 | 73 | 0.017 | (>37%) | 618 | 0.145 | |
| Delta1a | 10,617,271 | 72 | 0.007 | (>30%) | 375 | 0.035 | (>65%) |
| Delta1b | 9,888,100 | 57 | 0.006 | (>50%) | 624 | 0.063 | (>65%) |
| Delta4 | 5,465,468 | 177 | 0.032 | (>60%) | 675 | 0.124 | (>50%) |
| Spirochete | 2,393,688 | 99 | 0.041 | (>5%) | 88 | 0.037 | (>63%) |

*SNP sites were filtered with a minimum coverage of 5× per site in each sample. When no SNP site was found due to very low or lacking reads from a symbiont, these individuals were excluded, using a cut-off based on lateral coverages (i.e. % reference genetic sites ≥5× covered by reads) as indicated in brackets.

**SNP sites were filtered with a SNP p-value cut-off of 0.01, genetic sites within the 5~95 percentile range based on total read-depth distribution, a minimum read-coverage of 1× per site in each sample, and a minimum minor allele frequency of 0.01. When no SNP site was found due to very low or lacking reads from a symbiont, these individuals were excluded using a cut-off based on lateral coverages (i.e. % reference genetic sites covered by reads) as indicated in brackets.

**Supplementary Table S5** Statistical comparisons of pairwise genetic distances for the host mitochondria and symbionts, among three categories; (i) within the same combination of location plus A- or B-hosts ("Within"), (ii) between A- and B-hosts ("Between A and B"), and (iii) between locations. See the caption for Figure 4 in the main article for the details of the comparisons.

| Kruskal-Wallis test | | | Post-hoc Dunn tests | | |
|---|---|---|---|---|---|
| **Subject** | **$\chi 2$** | **p-value** | **Compared categories** | **z** | **p-value** |
| **a**; Mitochondria | 85.77 | 2.2E-16** | Within vs. Between A and B | 8.98 | 7.9E-19** |
| | | | Within vs. Between locations | 2.53 | 1.1E-02* |
| | | | Between A and B vs. Between locations | 6.45 | 1.7E-10** |
| **b**; *Ca.* Thiosymbion | 79.72 | 2.2E-16** | Within vs. Between A and B | 8.02 | 3.1E-15** |
| | | | Within vs. Between locations | 0.62 | 0.54 |
| | | | Between A and B vs. Between locations | 7.41 | 2.2E-13** |
| **c**; Gamma3 | 30.73 | 2.1E-07** | Within vs. Between A and B | 3.24 | 1.8E-03** |
| | | | Within vs. Between locations | 5.52 | 1.0E-07** |
| | | | Between A and B vs. Between locations | -2.28 | 2.3E-02* |
| **d**; Delta1a | 1.97 | 0.37 | - | | |
| **e**; Delta1b | 10.41 | 1.5E-2* | Within vs. Between A and B (in Cavoli) | 0.53 | 0.71 |
| | | | Within vs. Between locations (in B-hosts) | 2.84 | 2.7E-2* |
| | | | Between A and B vs. Between locations | -0.12 | 0.91 |
| **f**; Delta4 | 50.6 | 1.0E-11** | Within vs. Between A and B | 1.56 | 0.12 |
| | | | Within vs. Between locations | 6.83 | 2.6E-11** |
| | | | Between A and B vs. Between locations | -5.26 | 2.2E-07** |
| **g**; Spirochete | 2.03 | 0.36 | - | | |

(Asterisks denote statistical significance of p-value < 0.05 (*) and p-value < 0.01 (**)).

**Supplementary Table S6** Statistical tests on genetic co-divergence patterns of host mitochondria and symbionts; (**a**) Mantel tests on the correlation between pairwise genetic distance of mitochondria and pairwise genetic distance of each symbiont. (**b**) Kendall's rank correlation test on the correlation between the relative symbiont abundance (as shown in Figure 2) and Mantel's R (as shown in (**a**)).

**a**; Mantel tests on genetic co-divergence (host mitochondria and each symbiont)

| Symbiont | Mantel's R | p-value |
|---|---|---|
| *Ca*. Thiosymbion | 0.988 | 0.001** |
| Gamma3 | 0.230 | 0.001** |
| Delta1a | 0.145 | 0.048* |
| Delta1b | 0.283 | 0.002** |
| Delta4 | 0.060 | 0.017* |
| Spirochete | 0.015 | 0.349 |

**b**; Kendall's rank correlation test (Mantel's R and relative abundance)

| Kendall's rank correlation tau | z | p-value |
|---|---|---|
| 0.354 | 10.65 | 2.2E-16** |
